# Across kingdom biased CYP-mediated metabolism via small-molecule ligands docking on P450 oxidoreductase

**DOI:** 10.1101/2020.09.23.310391

**Authors:** Simon Bo Jensen, Sara Thodberg, Shaheena Parween, Matias E. Moses, Cecilie C. Hansen, Johannes Thomsen, Magnus B. Sletfjerding, Camilla Knudsen, Rita Del Giudice, Philip M. Lund, Patricia R. Castaño, Yanet G. Bustamante, Flemming S. Jørgensen, Amit V. Pandey, Tomas Laursen, Birger Lindberg Møller, Nikos S. Hatzakis

## Abstract

Metabolic control is mediated by the dynamic assemblies and function of multiple redox enzymes. A key element in these assemblies, the P450 oxidoreductase (POR), donates electrons and selectively activates numerous (>50 in humans and >300 in plants) cytochromes P450 (CYPs) controlling metabolism of drugs, steroids and xenobiotics in humans and natural product biosynthesis in plants. The mechanisms underlying POR-mediated CYP metabolism remain poorly understood and to date no ligand binding has been described to regulate the specificity of POR. Here, using a combination of computational modeling and functional assays, we identified ligands that dock on POR and bias its specificity towards CYP redox partners. Single molecule FRET studies revealed ligand docking to alter POR conformational sampling, which resulted in biased activation of metabolic cascades in whole cell assays. We propose the model of *biased metabolism*, a mechanism akin to biased signaling of GPCRs, where ligand docking on POR stabilizes different conformational states that are linked to distinct metabolic outcomes. Biased metabolism may allow designing pathway-specific therapeutics or personalized food suppressing undesired, disease related, metabolic pathways.

## MAIN TEXT

Across the kingdoms, metabolic control is mediated by the dynamic assemblies and function of multiple redox protein partners with cytochromes P450 (CYPs) and NADPH-dependent cytochrome P450 oxidoreductase (POR) as key elements ^1–3^. POR transfers electrons to the heme iron of CYPs and other redox partners selectively activating them ^4–10^. In plants, the coordinated assembly of POR-CYP complexes in dynamic metabolons enables on demand *in vivo* production of natural products to fend off, counteract or adapt to biotic or abiotic environmental stress, as recently demonstrated in the crop plant sorghum ^1,11^. Metabolon formation also regulates parts of primary metabolism. In humans, POR-CYP assemblies serve to properly balance the metabolism of drugs, steroids, fatty acids, xenobiotics, and bio-active plant natural products in foods ^2,4^. Mutations in human POR alter POR specificity towards activation of CYPs leading to severe disorders with multiple clinical manifestations varying from skeletal malformations with craniosynostosis (similar to Antley-Bixler Syndrome) to ambiguous genitalia and disorder of sexual development, amongst others ^3,4,12–14^. Exploiting this regulatory layer is central for the treatment of metabolic disorder and tailored biosynthesis of natural products, however the mechanisms regulating, or biasing, POR specificity towards CYPs are not well understood.

Biased specificity has historically been observed to underlie function of signaling hubs like the G protein-coupled receptors (GPCRs). Docking of structurally diverse ligands biases GPCR conformational sampling, stabilizing distinct conformational states and thus the corresponding signaling pathways, a phenomenon called biased agonism ^15–17^. POR acts as a metabolic hub: Its conformational sampling and specificity towards CYPs is dependent on regulatory cues ^5,6,18^ and mutations ^4,19^ indicating that POR specificity of activating metabolic cascades may operate via mechanisms akin to biased agonism. However, to date no small molecules are known to target metabolic hubs like POR and allosterically control downstream metabolic pathways.

Here, we show that the specificity of POR towards diverse electron acceptors can be tuned by small molecules. We demonstrate using computational modeling that the small molecules serve as ligands docking on POR. Comparative *in vitro* activity assays on a set of diverse electron acceptors display that the ligands bias the specificity of both human and plant POR rather than inhibiting their function. Single molecule Förster Resonance Energy Transfer (smFRET) provides mechanistic insights showing that ligand docking biases conformational sampling of plant POR, providing a link from biased conformational sampling to biased redox partner specificity. Lastly, we show that ligands alter CYP-mediated steroid hormone metabolism in human cells and microsomes emphasizing the biological relevance and applicability of controlling metabolic outcomes by targeting POR. Our data support a model of *biased metabolism*, a mechanism akin to biased signaling of GPCRs: POR conformational states are optimized to interact with certain CYPs and are linked to distinct downstream metabolic outcomes. Ligand-mediated control of POR conformational sampling thus appears to inhibit the activation of a subset of CYPs and/or enhance activation of others offering a new paradigm of metabolic control.

### Docking of POR ligands induce biased specificity towards electron acceptors

Using computational docking simulations on POR crystal structures, we assessed the possible docking of small-molecule ligands that are known to affect the activity of specific CYPs (Fig 1A). Based on docking simulations and functional activity assays, three structurally diverse ligands with promising effects on POR function were selected for detailed studies (Fig 1B). Tested molecules showing weaker or no effects on POR function are displayed in Supplementary Fig 1. The three promising candidates were; a) rifampicin, an antibiotic that induces expression and function of several CYPs ^20^, b) cyclophosphamide, a chemotherapeutic prodrug which induces expression of several CYPs ^21^, and c) dhurrin, a plant defense compound which biosynthesis requires the coordinated assembly and function of POR and CYPs in dynamic metabolons ^1,11^.

**Fig. 1.**
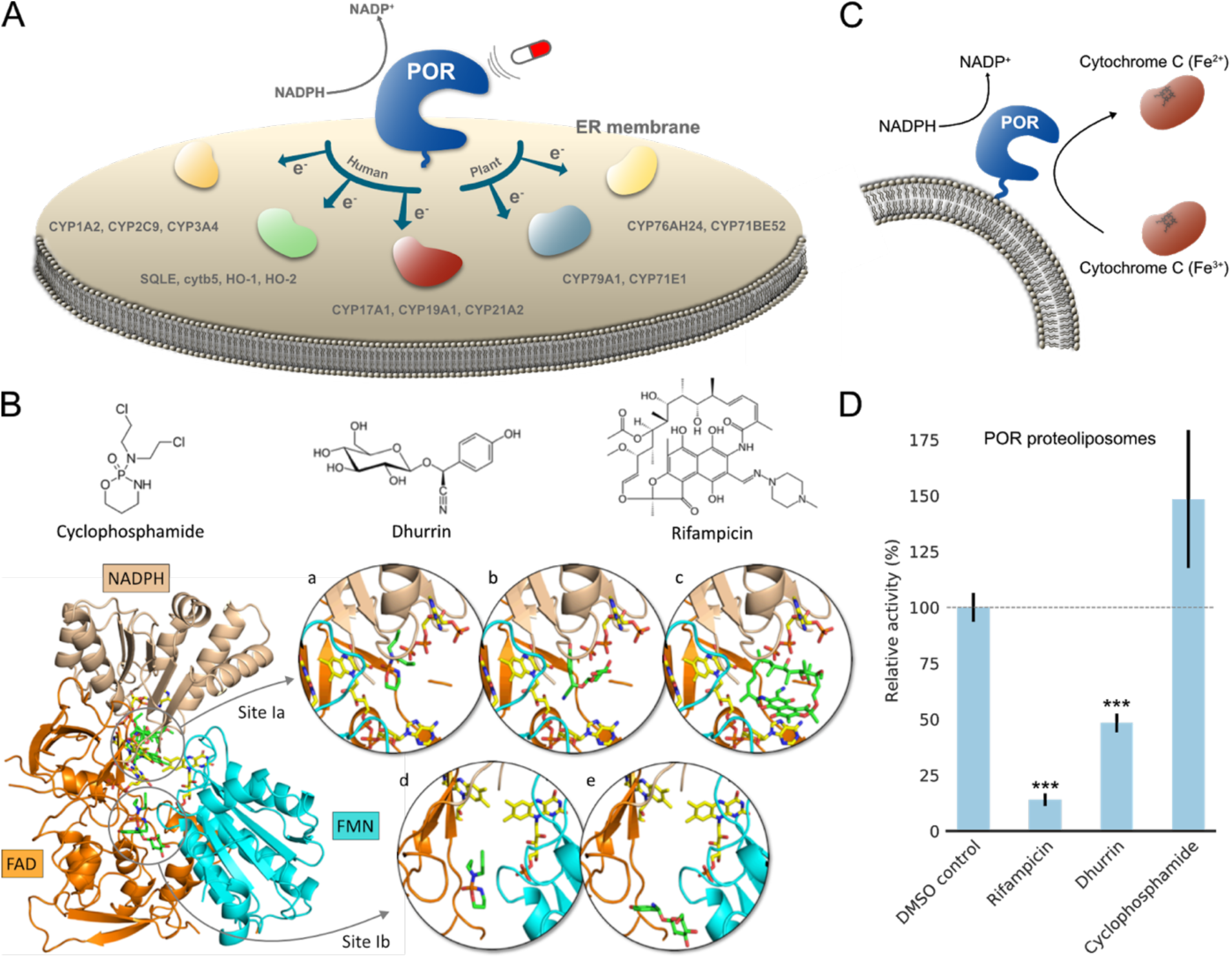
Small molecules dock on human POR and regulate electron transfer in vitro. A) POR is the omnipotent electron donor to all CYPs in the ER membrane, activating metabolic cascades in both human and plants by transferring electrons to redox partners. Targeting POR with small-molecule ligands may bias metabolic outcomes and regulate basic metabolism in humans or tune the formation of natural products in plants. B) Molecular structures of small-molecule ligands and their respective docking. Ligands (green) were docked on human POR with cofactors (yellow) in a compact conformation (PDB 3QE2) in Sites Ia and Ib determined from SiteMap analysis (see Supplementary Fig 2 and Supplementary Table 1). Insets display the predicted binding conformations of cyclophosphamide (a+d), dhurrin (b+e), and rifampicin (c). See Supplementary Fig 3 and Supplementary Table 1 for detailed interactions and binding energies. C) In vitro activity of human POR proteoliposomes measured by the commonly used Cytc assay ^23^. D) Ligands bias human POR capacity to reduce Cytc in proteoliposomes at 100 μM, acting either as agonist (cyclophosphamide) or inverse agonists (dhurrin and rifampicin). The bar plot represents the mean of at least three measurements normalized to DMSO controls (see supplementary methods for experimental details). Error bars represent ±SEM (* p<0.05; ** p<0.01; *** p<0.005).

Potential docking sites on POR were identified based on a SiteMap analysis ^22^ on human POR in a compact conformation (PDB 3QE2 ^13^) and rat POR in an extended conformation (PDB 3ES9 ^7^). The analysis on the two isoforms sharing 94% sequence identity yielded five potential docking sites on each structure (see Supplementary Fig 2 and Supplementary Table 1). The ligands were docked into these docking sites, and all displayed a clear preference towards Site I on both structures with estimated binding energies ranging from −5 to −7 kcal/mol (see Supplementary Table 1). On human POR, Site I extends throughout the interface between the FMN-, FAD- and NADPH-binding domains comprising a relatively large volume. Both cyclophosphamide and dhurrin docked into two subsites of human POR Site I, called Ia and Ib, with almost equal binding energies, while rifampicin only docked into Site Ia (see Fig 1B). Site Ia forms a cavity partly comprised by the FAD and NADPH cofactors with the docked ligands extensively exposed to the surrounding solvent, while Site Ib lies at the interface between the FMN- and FAD-binding domains distant from the cofactors and further embedded into the protein structure. On rat POR, ligands only docked into Site I which aligns with Site Ia of human POR. All three ligands were predicted to interact via H-bonds and pi-pi stacking to several amino acid residues on both POR isoforms (see Supplementary Figs 2-3 and Supplementary Table 1 for detailed interactions). Notably, amino acid residues G539 and R600 of human POR, both within a 5 Å distance from the docked ligands in Site Ia, are associated with POR deficiency. Patients with pathogenic mutations G539R and R600W suffer from disorder of sexual development due to low production of sex steroids indicating that this site is important for POR specificity towards CYPs ^4,12^.

**Fig. 2.**
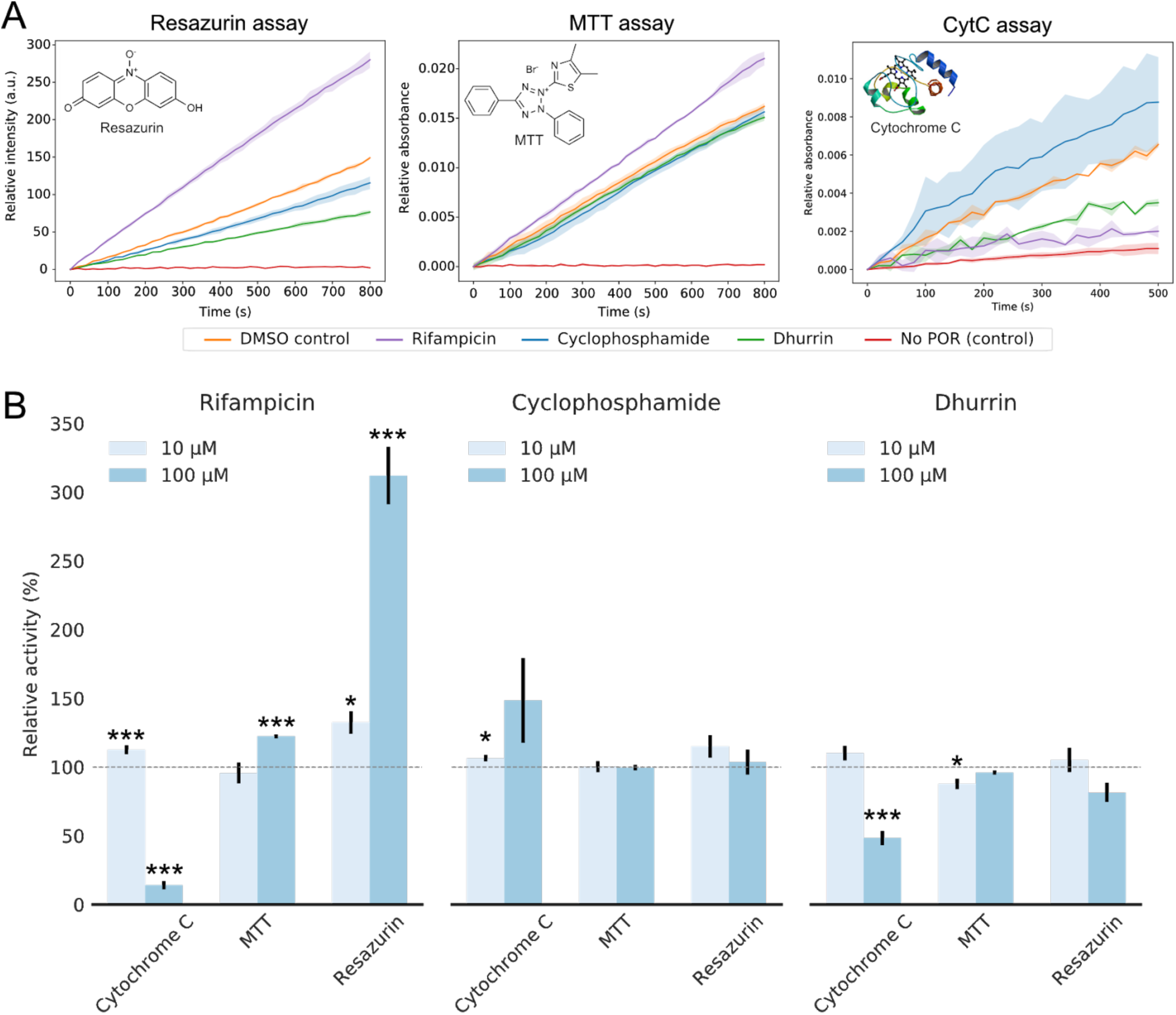
Small-molecule ligands bias specificity of human POR to reduce diverse electron acceptors. A) Human POR proteoliposome activity to reduce diverse electron acceptors was assessed using 100 μM NADPH and 10 μM RS (left), 500 μM MTT (middle) or 40 μM Cytc (right) by monitoring changes in absorbance (550 nm for Cytc, 610 nm for MTT) or fluorescence (582 nm for RS). Note the increased noise due to less sensitive UV-VIS readout for Cytc. All activity traces depict the average ± SD of at least three independent measurements. POR activity was extracted by fitting the linear region of the traces. B) Ligands affect the electron donating capacity of human POR differentially dependent on the electron acceptor indicating biased specificity. Rifampicin reduces POR activity towards Cytc, has a small effect on MTT reduction and enhances POR activity to reduce the electron acceptor resazurin by 3-fold. Cyclophosphamide results in minute increased activity towards Cytc, while dhurrin reduces activity towards Cytc. The bar plot represents the mean of at least three measurements normalized to DMSO controls with propagated error (see supplementary methods for details). Bar chart error bars represent ±SEM (* p<0.05; ** p<0.01; *** p<0.005).

We evaluated whether ligand docking predicted by the simulations affects POR function by performing *in vitro* functional assays on human POR proteoliposomes using cytochrome *c* (Cyt*c*) as an electron acceptor (Fig 1C-D, see Supplementary methods and Supplementary Fig 4 for spectral controls). All three ligands displayed a strong effect on the capacity of POR to reduce Cyt*c*. Rifampicin caused an activity decrease to 14±3% of control, defined as POR activity without drug in otherwise identical conditions. Dhurrin caused a decrease to 48±5% of control, while cyclophosphamide appeared to cause an increase to 149±31% of control. All compounds were tested at 100 μM using 40 μM Cyt*c* and 100 μM NADPH as substrates. The fact that the docked ligands modulate activity, supports that POR can be a target for metabolic regulation and a modulator of therapeutic activities. Furthermore, it shows that direct docking on POR should be taken into consideration when screening for drug-CYP interactions.

**Fig. 3.**
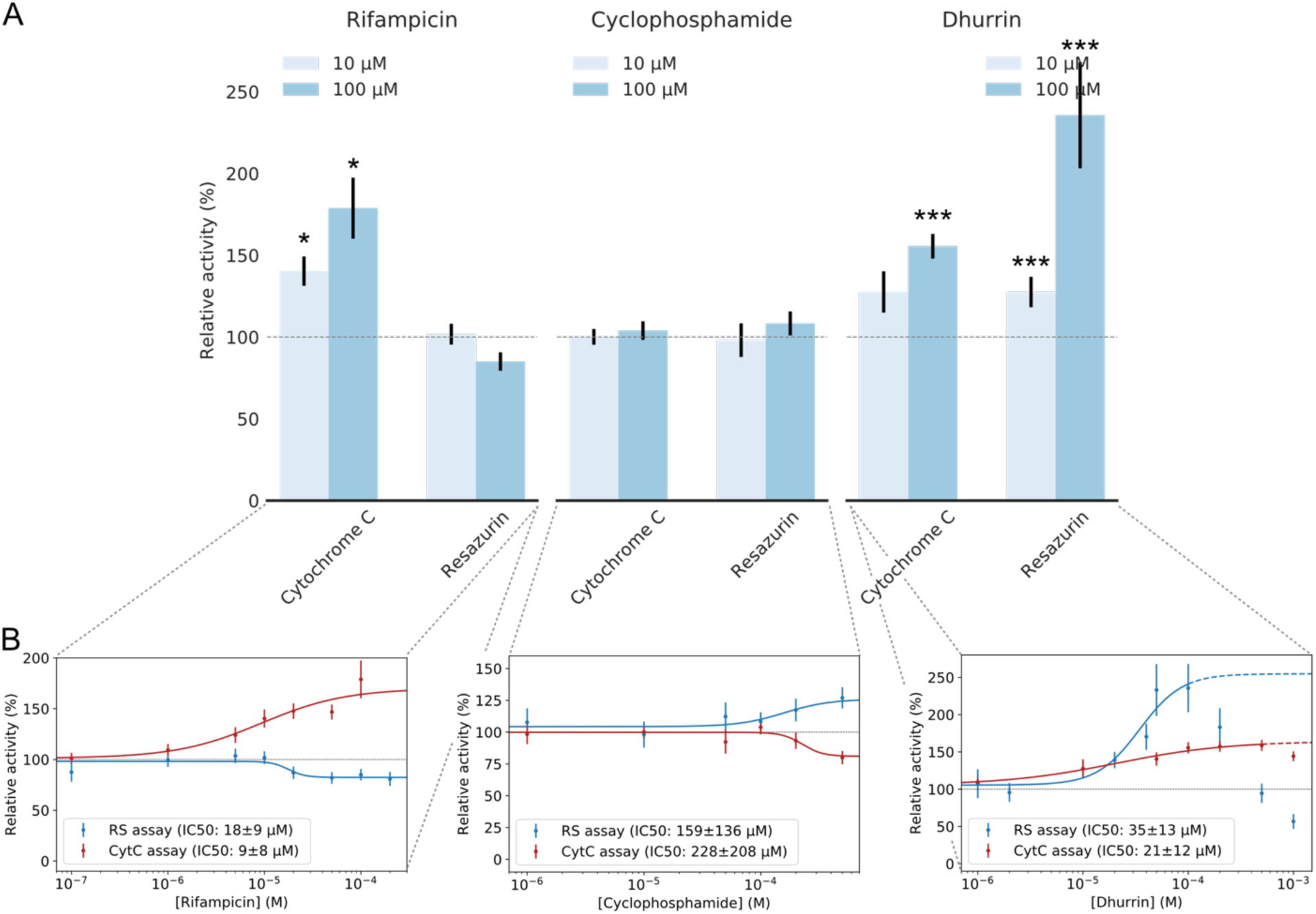
Small-molecule ligands bias specificity of plant POR (SbPOR2b) to reduce diverse electron acceptors. A) Effects of small-molecule ligands on SbPOR2b activity in proteoliposomes using Cytc and RS as electron acceptors. B) Dose-response curves of rifampicin, cyclophosphamide and dhurrin in the Cytc and RS assays, respectively. Rifampicin acts as an agonist towards Cytc enhancing its reduction rate and inverse agonist towards RS reducing the reduction rate. Cyclophosphamide displays the reverse effect acting as an inverse agonist towards Cytc reduction and agonist towards RS reduction. Dhurrin acts as an agonist towards both Cytc and RS reduction at low micromolar concentrations. The fact that ligands display differential effects on SbPOR2b activity to reduce the two electron acceptors indicates biased specificity of POR. IC50 values are extracted from the Hill equation. The bar plots and dose-response curves represent the mean of at least three independent measurements normalized to controls with propagated error (See Supplementary Fig 5 for raw data). Error bars represent ±SEM (* p<0.05; ** p<0.01; *** p<0.005).

**Fig. 4.**
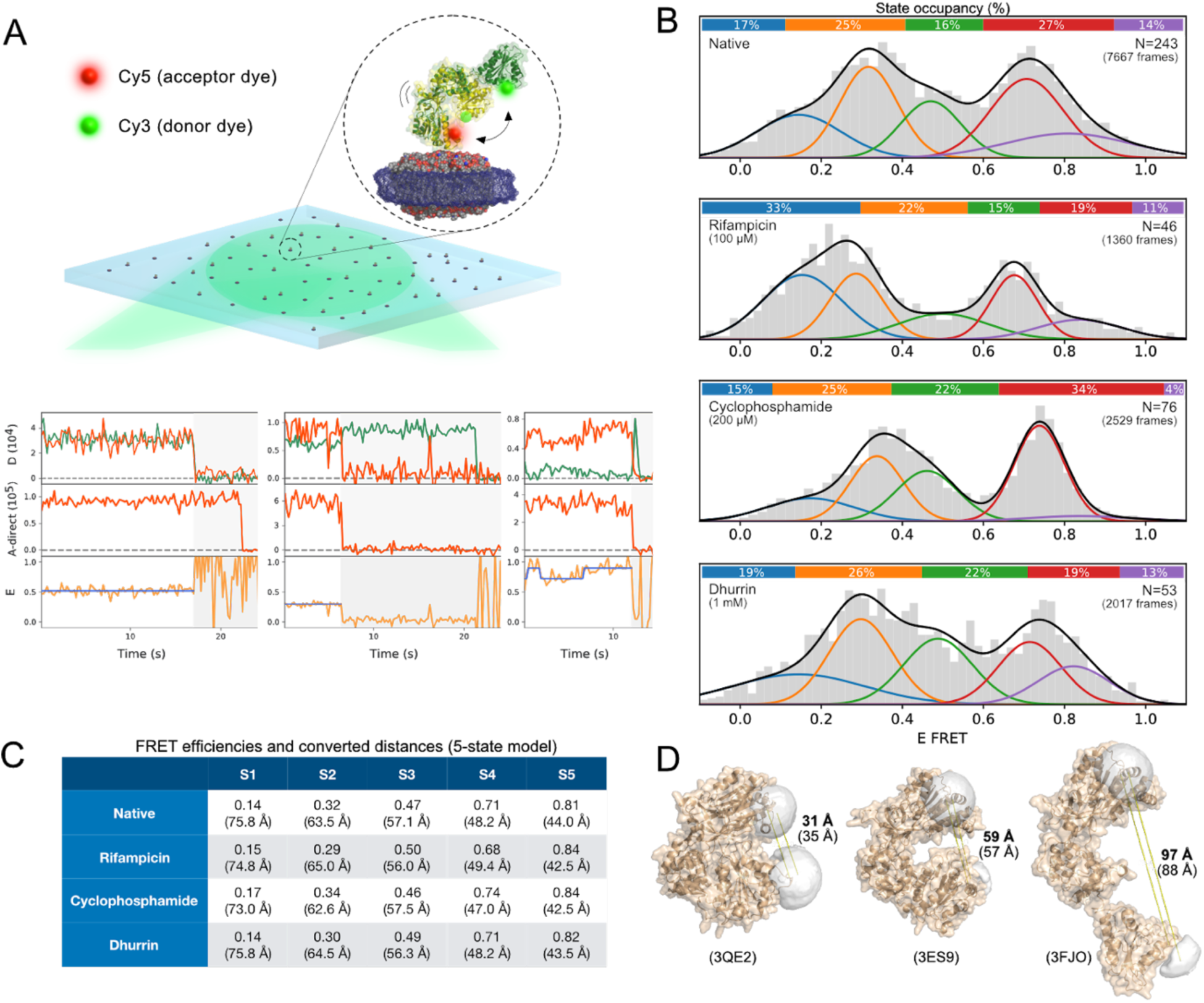
Direct observation of POR biased conformational sampling by small-molecule ligands using smFRET. A) Illustration of smFRET assay using TIRF microscopy. Top; SbPOR2b is site-specifically labeled with Cy3/Cy5 fluorophores, reconstituted in lipid nanodiscs and tethered on a passivated microscope surface. Bottom; representative smFRET traces displaying FRET states and dynamic transitions between them (see Supplementary Fig 7 for more examples). B) Distribution of FRET efficiencies in the absence and presence of ligands. Distributions are optimally fit with 5 states for all conditions as determined from BIC (see Supplementary methods and Supplementary Fig 8) with average distances ranging from ∼40 to ∼80 Å. Rifampicin, cyclophosphamide and dhurrin alter the occupancies of each of the five FRET states indicating biased conformational sampling. Colored bars on top of histograms represent occupancies of each state. C) FRET efficiencies and converted inter-dye distances obtained from five-state gaussian mixture models. D) Homology modeling of SbPOR2b from crystal structures of POR isoforms (3QE2, 3ES9 and 3FJO) with simulated inter-dye distances (bold) and Cα-Cα distances (brackets).

Given that point-mutations in human POR can lead to altered specificity towards CYP isoforms 4,19, we tested whether small-molecule ligands can bias the specificity of human POR to reduce diverse electron acceptors. Comparative *in vitro* activity assays were carried out using commonly employed artificial electron acceptors of POR; Cyt*c* ^23,24^, resazurin (RS) ^25^ or 3-(4,5-dimethylthiazol-2-yl)-2,5-diphenyltetrazolium bromide (MTT) ^26^ (Fig 2A). Each of the assays relies on spectral changes of the electron acceptor upon reduction by POR and thus directly reports on POR activity towards reducing the specific redox partner (see Supplementary methods for details). Cyclophosphamide appeared to cause an increase in POR capacity to reduce Cyt*c* (149±31% of control) but had no significant effect in neither the MTT nor RS assay when tested at 10 or 100 μM (Fig 2B). This supports that binding close to FAD and NADPH cofactors does not *apriori* reduce or eliminate activity. Dhurrin, which caused an activity decrease in the Cyt*c* assay (48±5% of control), only showed minor effects in the MTT and RS assays (96±2% and 82±7% of control, respectively). The presence of rifampicin caused a dramatic increase in POR capacity to reduce RS (312±21% of control) and a smaller but significant increase in reduction of MTT (122±2% of control; see Fig 2B). This is striking as rifampicin decreased POR capacity to reduce Cyt*c* (14±3% of control at 100 μM) and highlights that small-molecule ligands can bias the specificity of POR towards reducing diverse electron acceptors in a way similar to biased agonism of GPCRs.

### POR ligand binding and biased specificity pertain across kingdoms

POR plays an omnipotent role as an electron donor to microsomal CYPs in all eukaryotes and serves as a key metabolic hub in plants as well as humans ^1^. To test whether the effect of small-molecule ligand binding to POR is an omnipotent phenomenon underlying the regulation of POR in organisms from different kingdoms, we performed dose-response experiments on POR2b from the crop plant *Sorghum bicolor* (*Sb*POR2b) in proteoliposomes (Fig 3A+B). Rifampicin increased *Sb*POR2b capacity to reduce Cyt*c* while reducing the capacity in the RS assay (179±19% and 85±6% of control at 100 μM, respectively) with IC50 values of 9±8 μM and 18±9 μM, respectively (see Fig 3B and Supplementary Fig 5 discussing an observed lag phase). Cyclophosphamide caused the opposite effect resulting in an activity decrease in Cyt*c* reduction and increase in RS reduction (80±5% and 127±8% of control at 500 μM, respectively) with IC50 values of 228±208 μM and 159±136 μM, respectively. Note the large standard deviations due to low affinities. Careful inspection of the rifampicin and cyclophosphamide IC50 curves is reminiscent of biased agonism (Fig 3B). Rifampicin appears to operate as an inverse agonist inhibiting RS reduction and agonist increasing Cyt*c* reduction. Cyclophosphamide, on the other hand, displays the opposite behavior and operates as an agonist towards RS reduction and inverse agonist towards Cyt*c* reduction. The fact that the specificity of both human and plant POR isoforms can be biased by the tested ligands, albeit to a different extend, probably due to a low sequence identity of 38%, indicate that biased agonism may not be an exclusive property of receptor-mediated signaling ^15^ but also a method to regulate the function of metabolic cascades across kingdoms.

**Fig. 5.**
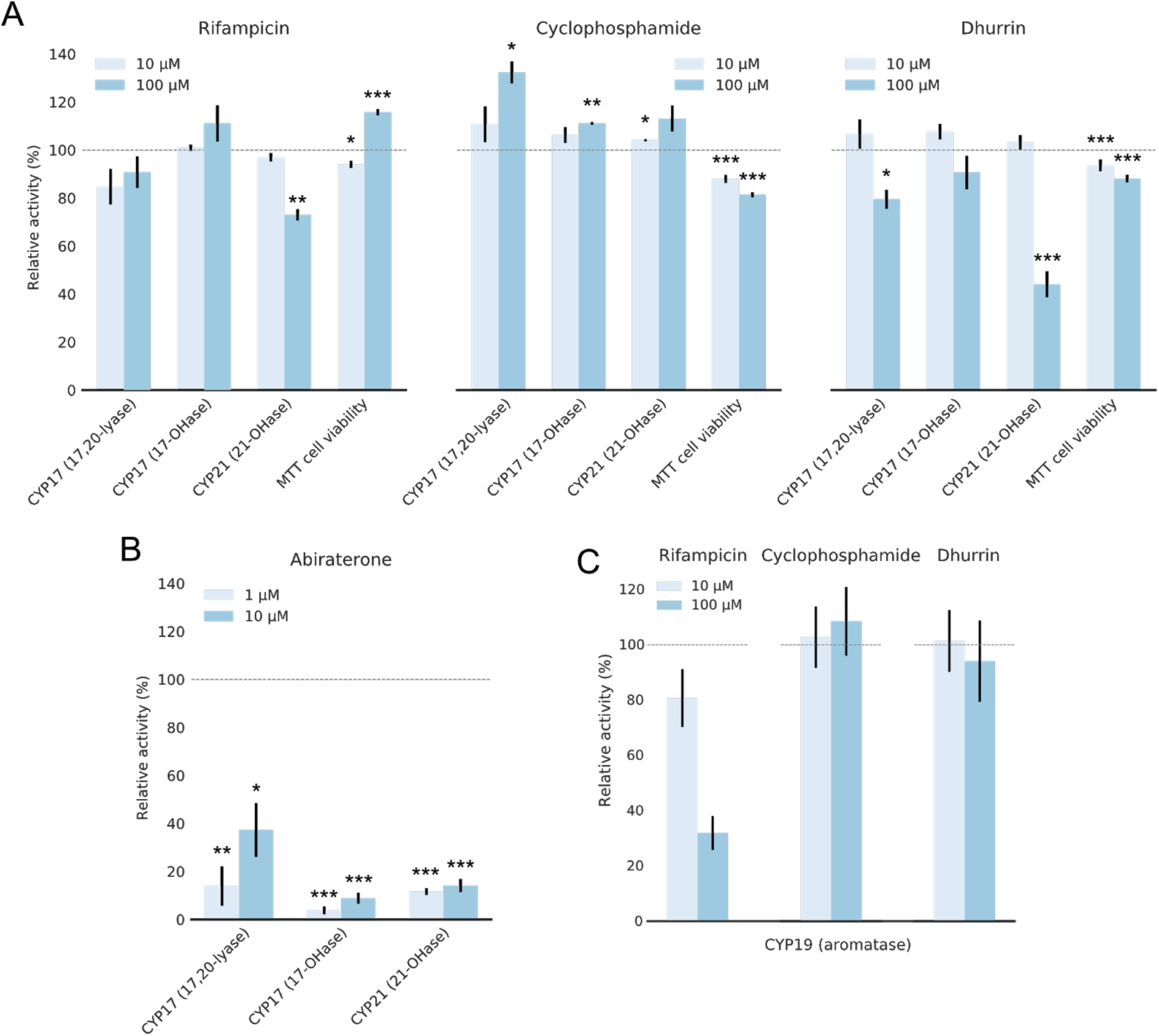
Biased Metabolism: Small-molecule ligands bias steroidogenic CYP-activities in human cells and microsomes. A) A human adrenocortical cell line (NCI-H295R) was used to assess the effect of small-molecule ligands on steroidogenic CYP17A1 and CYP21A2 hydroxylase activity, and CYP17A1 lyase activity, using radiolabeled substrates. Cell viability was assessed based on MTT reduction. Rifampicin shows a small inhibiting effect towards CYP21A2. The cells display increased MTT reduction indicating increased reductase activity. No significant effects on CYP17A1 activities are observed. Cyclophosphamide causes a small increase in both CYP17A1 and 21A2 activities, while MTT reduction decreases slightly. Dhurrin causes inhibition of both CYP17A1 and CYP21A2 activities. Interestingly, 17,20-lyase activity is affected more significantly than 17-OHase activity. B) Abiraterone was used as a control inhibitor of CYP17A1 and CYP21A2 in H295R cells. C) The effect of ligands on CYP19A1 activity was assessed on microsomes from a human choriocarcinoma cell line (JEG3). Rifampicin appears to show a concentration dependent inhibitory effect on CYP19A1 activity (32±6 % of control). Cyclophosphamide and dhurrin display no significant effect. A-C) Error bars represent ±SEM of at least three biological replicates normalized to DMSO controls with propagated error (* p<0.05; ** p<0.01; *** p<0.005; see Supplementary Fig 9 for raw data).

The natural product dhurrin caused increased *Sb*POR2b activity towards both Cyt*c* and RS (156±8% and 236±32% of control at 100 μM, respectively) with IC50 values of 21±12 μM and 35±13 μM, respectively. The observed effect is inverted at concentrations above 100-500 μM dependent on the assay, indicating a negative feedback loop type of mechanism downregulating dhurrin production in plants at high dhurrin concentrations (see Supplementary methods and Supplementary Fig 4 for control on dye photophysics). This may originate from lower docking affinity of dhurrin to alternative docking sites (see Supplementary Fig 2 and Supplementary Table 1) and can be studied in the future. Ligands also have an effect on POR reconstituted in detergent micelles (Supplementary Fig 6). Although the amplitude and effects are different in detergent compared to liposomes, it highlights that the observed effects are not induced by altered properties of the lipid bilayer. The fact that each ligand introduces diverse effects on POR capacity to reduce electron acceptors indicates that POR operates as a central metabolic hub integrating multiple layers of regulatory inputs (ligand conditions, subcellular localization, membrane environment and mutations) to tune the transferred electrons to CYPs consequently controlling metabolic cascades. This opens up the possibility to use POR as a target for regulating natural product biosynthesis in plants, basic metabolism in humans, and optimize synthetic biology approaches for production of bioactive metabolites ^1,11^.

### Direct observation of POR conformational sampling and its remodeling by ligands

Biased specificity is well established for receptors, and documented to operate via biased conformational sampling ^15–17^. POR is a highly dynamic protein oscillating between compact and extended conformations to execute electron transfer to CYPs. This has been verified by ensemble techniques including electron paramagnetic resonance (EPR) ^10^, nuclear magnetic resonance (NMR) ^6,9^, small-angle X-ray scattering (SAXS) ^6^, small-angle neutron scattering (SANS) ^5^, fluorescence ^24^, and stopped-flow ultraviolet-visible (UV-VIS) spectroscopy ^27^, providing insights into POR conformational sampling recently confirmed by smFRET burst analysis ^18,28^. Mutations known to control POR specificity and to cause metabolic disorders are often found in the hinge region of POR that controls conformational dynamics ^4,12,13^. We therefore hypothesized that POR biased specificity might originate from biased conformational sampling.

We used Total Internal Reflection Fluorescence (TIRF) microscopy ^29–31^ to record smFRET traces and directly observe conformational sampling of *Sb*POR2b and its remodeling by ligands. Data were recorded using the Alternating-Laser Excitation (ALEX) methodology ^32^ that we and others have been using extensively ^28,29,33^. POR was site-specifically labelled with Cy3 and Cy5 fluorophores using a minimal cysteine full-length *Sb*POR2b variant with two solvent accessible cysteines (N181C/C536S/A552C) that we have recently used for smFRET without impairing activity ^18^. Dual-labeled *Sb*POR2b was reconstituted in nanodiscs, which maintain the native structure and minimize non-specific interactions with the microscope surface ^18,25^ (Fig 4A and Supplementary Fig 7 for raw images). By monitoring FRET of hundreds of single POR enzymes in parallel, we were able to quantify the conformational sampling of POR and its remodeling by the ligands (Fig 4A). A wide range of conformations with average FRET distances varying from ∼40 to ∼90 Å were observed (see Supplementary methods, Supplementary Fig 7 for representative traces and Supplementary Fig 8 for calibration using ALEX). Inspection of individual traces revealed relatively stable fluorescence and only rare transitions between FRET states (Fig 4A and Supplementary 7). This is expected as POR dynamics related to function takes place at the low millisecond time scale ^6,25,27^. Imaging with a temporal resolution of 200 ms thus results in FRET states representing the equilibrium between one or more protein conformations. Indeed, decreasing the temporal resolution from 200 ms to 1 s still results in multiple distinct FRET states, however with a slightly higher fraction of traces showing dynamic transitions (from 4-8% to 10-14%; see Supplementary Fig 8) due to longer observation times. Increasing temporal resolution to timescales faster than 200 ms was not possible without compromising signal-to-noise. Fluorescence cross correlation studies yield Pearson coefficients centered around zero for all conditions, indicating that transitions between conformational states are masked due to dynamics faster than the temporal resolution (200 ms), in agreement with our simulations and similar readouts for GPCRs ^16^ (Supplementary Fig 8). Thus, the observation of multiple discrete FRET states, as well as a low fraction of transitions between them, originates from long-lived protein states as previously observed ^34–37^.

The FRET distribution of the native enzyme (n=243) as well as all data combined (n=418) were best fit with a mixture of five gaussians implying at least five underlying FRET states (see BIC analysis in Supplementary methods and Supplementary Fig 8) in agreement with earlier studies ^18^. A five-state model was used to quantify the abundance of FRET states (Fig 4B-C). In the native form, the inter-dye distances of the five FRET states were 76 Å, 64 Å, 57 Å, 48 Å and 44 Å. We modeled the structure based on human POR in a compact conformation (PDB 3QE2 ^13^), rat POR in an intermediate conformation (PDB 3ES9 ^7^) and a human-yeast chimera in a fully extended conformation (PDB 3FJO ^38^) as no crystal structure exists of *Sb*POR2b (see Fig 4D and Supplementary methods). The expected inter-dye distances calculated from dye-linker Monte Carlo simulations ^39^ on the three homology models were 31 Å, 59 Å and 97 Å, respectively, in agreement with our earlier burst analysis studies ^18^, further supporting each FRET state reflects an equilibrium between multiple conformations. We call these equilibrium states S1-S5, respectively. The occupancies of the five states were 17%, 25%, 16%, 27% and 14%, respectively, implying that states S2 and S4 are the most dominant.

To evaluate the effect of ligands on POR conformational sampling, we measured smFRET on POR exposed to ligand concentrations well above IC50. Ligand docking did not significantly affect the center of the FRET states (gaussian means), but rather their occupancies, indicating a changed equilibrium between long-lived protein equilibrium states (Fig 4B-C). Rifampicin shifted the equilibrium towards S1 at the expense of S4 and S5. S1 practically doubles its occupancy from 17% to 33%, while S4 decreases from 27% to 19%, but also slightly shifts its center indicating S5 slightly decreases from 14% to 11%. Cyclophosphamide shifted the equilibrium towards S3 (22% from 16%) and S4 (34% from 27%) at the expense of S5 and S1 (4% from 14% and 15% from 17%, respectively). Dhurrin at 1 mM resulted in a small shift in the equilibrium between S3 and S4 (S3 increases from 16% to 22% while S4 decreases from 27% to 19%). Rifampicin and cyclophosphamide seem to have opposed effects on conformational sampling. Rifampicin shifts the equilibrium towards extended states whereas cyclophosphamide shifts the equilibrium towards intermediate and compact states (albeit not fully compact S5). Interestingly, rifampicin and cyclophosphamide also have opposing effects on *Sb*POR2b specificity when monitored *in vitro*. Rifampicin operates as an agonist on Cyt*c* reduction and inverse agonist on RS reduction, while the effect of cyclophosphamide is reversed. These data thus support a correlation between conformational sampling and substrate specificity. Ligand docking on POR appears to bias conformational sampling stabilizing certain equilibrium states at the expense of others, consequently promoting the inhibition or activation of a subset of electron acceptors.

### POR ligands bias steroid hormone metabolism in human cells and microsomes

We tested the efficiency of the three ligands to elicit a physiological response on steroidogenic CYP activities in cells (Fig 5A, see Supplementary methods). Using a human adrenocortical cell line (NCI-H295R) we tested the effect of ligands on CYP17A1 and CYP21A2 ^40,41^ activities at 10 and 100 μM concentrations (Fig 5A). Using radiolabeled substrates, we were able to quantify the steroid hormone production of CYP17A1 and CYP21A2 and its remodeling by POR ligands (see Supplementary methods for details). Control experiments with abiraterone at 10 and 100 μM show specific inhibition of the two CYPs (Fig 5B and Supplementary Fig 9). Rifampicin caused no significant effect on CYP17A1 activity, but reduced CYP21A2 activity at 100 μM (73±2% of control; see Fig 5A and Supplementary Fig 9). The cells capacity to reduce MTT increased significantly (116±1% of control at 100 μM). The fact that CYP21A2 activity was reduced indicates that the observed increase in MTT reduction is not attributed to increased overall reductase expression ^20^, but rather remodeling of activity. Cyclophosphamide enhanced both CYP17A1 hydroxylase and lyase activity at 100 μM (111±1% and 132±5% of control, respectively), while having a small yet statistically insignificant effect on CYP21A2 activity (113±5% of control). Dhurrin caused a decrease in CYP21A2 hydroxylase and CYP17A1 lyase activities (44±5% and 80±4% of control at 100 μM, respectively). Interestingly, CYP17A1 hydroxylase activity was not significantly affected by dhurrin (91±7% of control). Specific inhibition of CYP17A1 lyase activity and not hydroxylase activity, that is relevant in the treatment of prostate cancer and polycystic ovary syndrome ^40^, may be achieved by targeting POR as an alternative to targeting CYP17A1 directly. One may argue that the tested ligands may dock on additional proteins, CYPs or receptors, and induce convoluted physiological responses additional to what is reported here ^20,21^. Our combined docking simulations, functional data, and smFRET structural data, illustrate that the ligands also dock on POR and affect its specificity towards reducing diverse electron acceptors.

To test the effect of ligands on CYP19A1 aromatase activity, we used microsomes extracted from a human choriocarcinoma cell line (JEG3) ^41^. Radiolabeled substrate was used to quantify CYP19A1 activity and its remodeling by POR ligands (see Supplementary methods for details). Cyclophosphamide and dhurrin did not show any significant effects at 10 nor 100 μM, while rifampicin appears to reduce CYP19A1 activity at 100 μM (32±6% of control) (Fig 5C). The microsome results further confirm that the observed effects are caused by biased activities and not via altered protein expression levels. The fact that the small-molecule ligands affect activities of steroidogenic CYPs in cells and microsomes is a key finding confirming the biological relevance of POR controlling metabolic cascades. The data confirm our docking simulations, *in vitro* assays and structural dynamics studies by smFRET, and support that biased conformational sampling of POR induced by ligands results in altered specificity towards CYPs. Ligand docking on POR thus appears to inhibit the activation of a subset of CYPs and/or enhance activation of others. We assign the term *biased metabolism* to this phenomenon since the mechanism is akin to biased signaling of receptors. We propose that biased metabolism represents an extra layer of regulatory control guiding metabolic pathways in complex cellular environments.

## Discussion and Conclusions

Protein conformational sampling, the dynamic exploration of conformational space, governs all major aspects of protein behavior from folding to function. Protein conformational states are often found to elicit distinct functional outcomes ^34,42,43^. GPCRs are the prime example of this phenomenon, acting as key signaling hubs with several conformational states linked to distinct downstream cellular processes ^15–17^. The combined studies presented here substantiates earlier studies on POR ^18^ and point to POR as being at the center of key metabolic hubs regulating the activation of CYPs and therefore metabolic pathways ^1,3,4^. While our data do not distinguish between the swinging and rotating motion models of POR ^8^ they provide a correlation between the existence of POR equilibrium conformational states with distinct phenotypic metabolic outcomes.

Key advancement in our understanding of protein-ligand interactions has opened the possibility for the development of drugs that act on the same protein and selectively stabilize protein states, consequently controlling different cellular outcomes, a phenomenon described as biased agonism 15. Biased agonism is well studied, brought into practice and explicitly exploited to underpin the function of signaling hubs like GPCRs. The combined data on the three chosen ligands serve as proof of concept of the mechanism of *biased metabolism*, a mechanism similar to biased agonism of GPCRs, but for metabolic hubs like POR. While the working concentrations (10-100 μM) are rather high, they support the electron transfer of POR to respond to different ligands in a pluripotent way; each ligand docking on POR stabilizes a distinct equilibrium state that is linked to distinct downstream metabolic outcomes. Interestingly, even small variations in the conformational sampling equilibrium induced by ligand docking suffices for large variations in metabolic outcomes. Although these ligands may also dock on additional proteins ^20,21^ our combined *in vitro*, single molecule and *in vivo* data clearly support that their docking on POR can facilitate biased metabolism. They also highlight the significance of testing small-molecule drugs and metabolites for docking on POR during early drug discovery and high-throughput CYP screening assays ^2^. We propose that biased metabolism represents an extra, hitherto unrecognized, layer of regulation capable of controlling metabolic pathways in complex cellular environments. Targeting POR may serve as a way of controlling POR-CYP interactions and regulate CYP-mediated metabolic pathways. Our findings pave the way for the *in silico* design of biased ligands specifically designed to tightly dock in Site I, or additional sites, and antagonize detrimental metabolic pathways while stimulating beneficial downstream processes to the relative exclusion of others, with the possibility to control basic metabolism and alleviate metabolic disorder.

The fact that plant and human isoforms, with a sequence identity of 38%, both present responses to ligand docking may indicate the presence of evolutionary conserved *hotspots* serving to tune the specificity of POR. This is further supported by our findings that the plant defense compound dhurrin, which is ingested as part of foods, docks on POR and affects its dynamics, function and metabolic response in human cell lines. Since dhurrin is a product of a pathway composed of two sequential CYPs and a glucosyltransferase, these findings open up the exciting possibility of controlling biosynthetic metabolism via feedback loop mechanisms in the production of high-value natural products. The effects of dhurrin on human POR-mediated metabolism further provide a mechanistic clue explaining why many substances in food, beverages and dietary supplements may affect basic metabolism and induce food-drug interactions ^44,45^. Such insights may pave the way for the design of personalized, plant-based food targeting POR to alleviate metabolic disorder or dietary supplements composed of natural products to suppress specific undesired metabolic pathways associated with disease. We anticipate that harnessing the structural basis of conformational sampling in biased metabolism may offer the design of metabolic pathway-specific ligands. This could have direct implications in biomedicine for enhancing therapeutically relevant metabolic pathways. Quantitative single molecule structural and functional studies will be crucial in this endeavor of deciphering and controlling POR-mediated metabolism via biased ligands.

## Author contributions

SBJ performed all *in vitro* activity assays, smFRET assays and data analysis with the help of JT, PML, YGB and MBS. SBJ performed homology modeling, Monte Carlo simulations and simulation of smFRET traces for cross correlation studies with the help of JT and MBS. Protein expression, purification and liposome reconstitution was performed by ST, CCH, CK, RDG with help of SBJ and MEM. Labelling of *Sb*POR2b and reconstitution in nanodiscs was performed by SBJ, CCH, ST, MEM and TL. Cell assays were performed by SP and PRC with help of ST, MEM and SBJ. FSJ performed docking simulations. SBJ and NSH wrote the manuscript with inputs from all authors. NSH conceived the research initiative, had the overall strategic planning with help of BLM and together with BLM were responsible for the overall project supervision. All authors discussed and evaluated all data.

## Data and materials availability

All data is available in the main text or the supplementary materials. Raw data in the format of .csv or .tif files are available upon request.

