## Supplementary information for "Across kingdom biased CYP-mediated metabolism via small-molecule ligands docking on P450 oxidoreductase"

### Materials and methods
Chemicals and materials

All chemicals were of analytical grade and purchased from Sigma-Aldrich (Merck) unless otherwise stated. Phospholipids, 2-dilauroyl-sn-glycero-3-phosphocholine (DLPC) and 1,2-dilauroyl-sn-glycero-3-phosphoglycerol (DLPG), and 1,2-dipalmitoyl-sn-glycero-3-phosphoethanolamine-N-(cap biotinyl) (Biotinyl Cap PE) were purchased from Avanti Polar Lipids. Dhurrin was chemically synthesized as described previously (*1*, *4*). Bio-Beads SM-2 were purchased from Bio-Rad. 2'5'-ADP Sepharose size exclusion chromatography (SEC) column Superdex 200 HR 10/30 for high-resolution preparative separation were from GE Healthcare Life Sciences. Bottomless 6 channel (30µL) sticky slides were purchased from ibidi GmbH. PLL(20 kDa) grafted with PEG(2 kDa) (PLL-PEG) and PLL(20 kDa) grafted with PEG-Biotin (3.4 kDa) (PLL-PEG-biotin) was purchased from SuSoS AG. NeutrAvidin protein was purchased from Thermo Fisher Scientific.

Radiolabeled substrates [$^3$H]-pregnenolone, [$^3$H]-androstenedione, [$^3$H]-progesterone, [$^{14}$C]-progesterone were obtained from PerkinElmer and American Radiolabeled Chemicals Inc. Silica gel-coated aluminum backed TLC plates were purchased from Macherey-Nagel. The tritium screens used for the autoradiography were purchased from Fujifilm. Trilostane was extracted in absolute ethanol (EtOH) from tablets commercially available as Modrenal® (Bioenvision, NY, USA). CYP17A1 was obtained from Cypex Limited. Human adrenal carcinoma cell line (NCI-H295R) and human placental JEG3 cell line was purchased from American Type Culture Collection (ATCC: CRL-2128 and HTB-36TM, respectively).

Protein expression and purification

Full-length human wild-type POR (NCBI reference sequence: NP_000932.3) was expressed in *E. coli* BL21(DE3) according to protocols (*2*). After expression, the bacterial membrane was extracted and used directly for protein reconstitution in liposomes without further purification. We note, that *E. coli* does not express any membrane-bound reductases nor CYPs which would infer with our POR activity assays.

Full-length *Sorghum bicolor* POR2b (*Sb*POR2b; NCBI reference sequence: XP_002444097.1), cloned in pET52b was expressed in *E. coli* NiCo21 (DE3) cells (New England Biolabs). A 100 mL starter culture of terrific broth (TB) supplemented with 50 µg/ml ampicillin was grown overnight at 37°C, 220 rpm. The starter culture was diluted into 1 L of TB supplemented with ampicillin in a wide bottom flask and incubated at 37°C. Expression was induced at $OD_{600nm} = 0.6$ by addition of IPTG to a final concentration of 1mM. (-)-riboflavin at a final concentration of 1 µg/ml was also supplemented. Cells were harvested after expression for 6 hours at 25°C. *Sb*POR2b mutant N181C / C536S / A552C[11] in pET52b was expressed in *E. coli* HI-Control™ BL21DE3 strain (Lucigen) in 2*400 mL TB cultures in wide bottom flasks. 1 µg/ml FMN and 1 µg/ml FAD were supplemented at the beginning of expression. Cells were harvested after 18 hours expression at 20°C. The enzymes were purified from isolated membranes using 2′5′-ADP Sepharose affinity chromatography and anion exchange chromatography as described previously (*19*, *26*). During purification of *Sb*POR mutants, 100 µM FMN and FAD were included to ensure excess cofactors.

Protein reconstitution in liposomes

Liposomes were prepared using a DLPC/DLPG lipid mixture (3:1 ratio) dissolved in DMSO and placed in vacuum for ~4 hours to obtain dry lipid films. While kept on ice, lipid films were rehydrated with either purified protein or bacterial membrane extract in a solution containing 50 mM cholate to obtain final lipid:protein ratios of ~200 and protein concentrations of 2-20 µM. After 1 hour of incubation on a shaking platform at 5°C, biobeads (Bio-Beads SM-2) were added to the mixture to extract detergent molecules. Upon additional 2 hours of incubation, samples were centrifuged briefly to remove biobeads followed by centrifugation at 5000 RPM and 5°C for 10 minutes. Supernatant was collected and transferred to eppendorf tubes. All samples were flash frozen and stored at -80°C until further use.

*In vitro* POR activity assays

The activity of full-length POR reconstituted in either detergent micelles (20 mM cholate) or liposomes (DLPC/DLPG, 3:1 ratio) was assessed spectrophotometrically as described elsewhere (*24–27*). Using Cyt*c*, MTT or RS as electron acceptors the change in absorbance (550 nm for Cyt*c* and 610 nm for MTT) or emission (570 nm excitation, 585 nm emission for RS) was monitored as a function of time. Cyt*c* and RS concentrations were both close to $K_m$ of the given electron acceptor, while MTT and NADPH were in excess amounts. Concentrations were 40 µM Cyt*c*, 10 µM RS, 500 µM MTT and 100 µM NADPH unless otherwise stated. POR activity was extracted from the slope of the linear region of each trace. All measurements were repeated at least three times and subsequently normalized to control measurements. Fluorescence intensities of resorufin in the presence of rifampicin were corrected according to a linear calibration curve (see Supplementary Fig 4).

Cell lines and culture media

Cells were cultured according to established protocols (*41*, *42*). Human placental JEG3 cells were cultured in minimal essential medium (MEM) with Earle's salts (Thermo Fisher Scientific) supplemented with 10% fetal bovine serum, 1% L-glutamine (200 mM GIBCO), 1% penicillin (100 U/ml; GIBCO), and streptomycin (100 µg/mL; Thermo Fisher Scientific). Human adrenocortical NCI-H295R (NCI-H295R) cells were grown in DMEM/Ham's F-12 medium containing L-glutamine and 15 mM HEPES (Thermo Fisher Scientific) supplemented with 5% NU-I serum (Becton Dickinson), 0.1% insulin, transferrin, selenium (100 U/mL; Thermo Fisher Scientific), 1% penicillin (100 U/mL; Thermo Fisher Scientific), and streptomycin (100 µg/mL; GIBCO) and passage numbers during the experiments remained below 30.

Preparation of microsomes and CYP19A1 activity assay using JEG3 cells

Microsomes were prepared from JEG3 cells as described previously (*42*). Briefly, JEG3 cells were collected near confluency and washed with cold PBS. The cell suspension was then centrifuged at 1500 g for 5 min to pellet the cells. The cell pellet was suspended in 100 mM $Na_3PO_4$ (pH 7.4) containing 150 mM KCl, and the cells were lysed by sonication. Unbroken cells and mitochondria were pelleted by centrifugation at 14,000 g for 15 min at 4°C. Microsomes containing endoplasmic reticulum were collected by ultracentrifugation at 100,000 g for 90 min at 4°C and resuspended in 50 mM $K_3PO_4$ (pH 7.4) containing 20% glycerol.

CYP19A1 activity was measured by the release of tritiated water from radiolabeled substrates during aromatization as described previously (*42*). Briefly, 40 µg microsomes extracted from JEG3 cells were incubated with 50 nM [$1_{\beta}$-$^{3}$H(N)]-androstenedione (~20,000 cpm/reaction) in buffer (100 mM NaCl, 100 mM potassium-phosphate, pH 7.4) for 5 minutes at 37°C on a shaking platform. The reaction was initiated by adding 1 mM NADPH. After 1 hour of incubation at 37°C, the reaction was stopped by adding a mixture of 5% charcoal and 0.5% dextran. Samples were vortexed for 40 sec and centrifuged at 13,000 RPM for 5 min. Supernatant was collected and diluted in scintillation liquid (Rotiszint Universal Cocktail; Carl Roth GmbH) before counting [$^{3}$H]-radioactivity. All measurements were repeated at least three times and subsequently normalized to controls.

#### CYP17A1 and CYP21A2 activity assay in H295R cell line

Steroidogenic CYP17A1 and CYP21A2 activities in H295R cells were quantified as described previously (*41*, *42*). In brief, cells were plated in six-well plates and treated with small-molecule ligands dissolved in 0.1% DMSO and normal growth medium for 24 hours, or for 4 hours for rifampicin, to ensure protein expression levels are not affected (*21*). After incubation, 1 µM trilostane (a specific blocker of HSD3B) was added to the medium for 90 min followed by addition of 1 µM radiolabeled substrate ([$^{3}$H]-17α-OH-progesterone or [$^{3}$H]-pregnenolone; ~50,000 cpm). After 1 hour of reaction, steroids were extracted from cell supernatants and separated by thin layer chromatography (TLC) on silicagel (SIL G/$UV_{254}$) TLC plates (Macherey-Nagel, Oensingen, Switzerland). The steroids were visualized on a Fuji FLA-7000 PhosphorImager (Fujifilm, Dielsdorf, Switzerland) and quantified using Multi Gauge software (Fujifilm, Dielsdorf, Switzerland).

Steroid conversion was assessed as a percentage of incorporated radioactivity in relation to total radioactivity measured for the whole sample. The conversion of 17α-OH-progesterone to 11-deoxycorticosterone was used as a measure for CYP21A2 hydroxylase activity. The conversion of pregnenolone to 17α-OH-pregnenolone and dehydroepiandrostenedione (DHEA) was used as a measure for CYP17A1 hydroxylase activity, while the specific conversion of 17α-OH-pregnenolone to DHEA was used as a measure for 17,20-lyase activity. All measurements were repeated at least three times and subsequently normalized to controls.

#### MTT cell viability assay

MTT reduction was used to evaluate cell viability and simultaneously quantify reductase expression/activity of cells upon drug incubation as described elsewhere (*42*). In brief, 100 µL of cell solution containing approximately $3x10^{4}$ cells were placed in a 96-well plate at a concentration of upon 24-hour incubation with drugs. MTT was added in each well to a final concentration of 0.8 mg/mL. Absorbance was measured at 610 nm on a plate reader to quantify reduction of MTT. All measurements were done in triplicates and normalized to DMSO controls.

#### Homology modelling and simulation of dye-dye distances

Since no 3D structure of *Sb*POR2b is available, the structure was modelled using SWISS MODEL automated online server (*47*) based on human, yeast and rat POR isoforms as previously described (*19*). The compact conformation of *Sb*POR2b was modelled based on human POR (PDB 3QE2 (*15*), chain A) while the intermediate and fully extended conformations were based on rat POR (PDB 3ES9 (*8*), chain A) and a human-yeast chimera (PDB 3FJO (*39*)), respectively. The isoforms share 38%, 40% and 36% sequence identity to *Sb*POR2b, respectively.

To convert the modelled Cα-Cα distances between residues N181 and A552 to expected dye-dye distances as measured by smFRET we used a toolkit developed by Kalinin et al (*40*). The toolkit employs a geometric accessible volume (AV) algorithm to predict the spatial distribution of donor and acceptor dyes. Using an approximated linker length of 14.0 Å, linker width of 4.5 Å and dye radius of 3.5 Å, the simulated dye-dye distances and modelled Cα-Cα distances differ by 3.7-8.9 Å depending on protein conformation (see main Fig 4, panel D).

#### Computational docking simulations on POR structures

The small molecules were constructed in Maestro (v. 9.8, Schrodinger 2018-3 release, Schrödinger, LLC, New York, NY, 2014) and prepared for docking in the LigPrep module (*48*). Both neutral and charged forms were generated and used as input structures for the docking. The rifampicin structure was taken from a structure of rifampicin monooxygenase complexed with rifampicin (PDB 5KOX (*49*)). Prior to docking, rifampicin was subjected to the LigPrep module, which produced three different conformations. After a short energy minimization these structures were used as input structures for the docking.

Identification of potential binding sites was carried out using the SiteMap software (v. 2.6, Schrödinger, LLC) (*23*). All ligands were docked on the human POR crystal structure (PDB code 3QE2 (*15*), A-chain) as well as the rat POR crystal structure (PDB code 3ES9 (*8*), A-chain) representing the compact and extended POR conformations, respectively, using Glide (v. 5.8, Schrödinger, LLC) with default settings (van der Waals scaling factor and partial charge cut-off at 0.80 and 0.15, respectively; standard precision and flexible ligand sampling; no constraints) (*50*),(*51*). The stability of top scoring docking conformations were evaluated to be stable by short MD simulations (embedded in water box, 10 ns) in Desmond (v. 4.3, Schrödinger, LLC) (*52*). All docking results are displayed in Supplementary Table 1.

Protein labeling and nanodisc reconstitution for smFRET

*Sb*POR2b mutant N181C/C536S/A552C containing two solvent accessible cysteines was labeled with Cy3 and Cy5 maleimide mono-reactive dyes (GE Healthcare) as previously described (*19*). A stoichiometric ratio of 1:2 Cy3/Cy5 was used for optimal labeling conditions. Labeled protein was reconstituted in lipid nanodiscs comprising membrane scaffold protein MSP1E3D1 and a mixture of DLPC:DLPG:Biotinyl Cap PE:DiO (69.8:25:4:1.2 ratio) according to protocols described previously (*19*, *26*). Purification of nanodiscs was achieved by size exclusion chromatography (SEC) (flow rate: 0.5 mL/min) on a preparative HPLC (Shimadzu) equipped with a Superdex 200 Increase 10/300 GL column (Amersham Pharmacia Biotech; diameter 10 mm; length 300 mm) using a mobile phase of 50 mM Tris-HCl (pH 7.5), 100 mM NaCl. Elution of protein, Cy3 and Cy5 was continuously monitored by absorbance at 280, 550 and 650 nm, respectively. Selected fractions of nanodiscs containing POR were collected, flash frozen and stored at -80°C until further use.

Surface preparation and nanodisc immobilisation for smFRET

Dual-labeled POR reconstituted in lipid nanodiscs was immobilised on a PLL-PEG functionalized surface. The surfaces were prepared according to previously described protocols (*26*, *30*). In brief, glass coverslips were cleaned, dried under nitrogen flow and plasma etched (Harrick Plasma Cleaner PDC-32G-2) for 5-10 min at 60 Pa. Immediately after plasma etching, flow chambers were assembled using 6-channel sticky slides (Ibidi) and each chamber was incubated with a 1:100 mixture of PLL-PEG-biotin/PLL-PEG in HEPES buffer (pH 5.6) for at least 1 hour. Chambers were flushed with 1 mL buffer to remove excess PLL-PEG and afterwards incubated with 0.1 g/L NeutrAvidin for at least 10 min. The chambers were stored at 5°C until further use. Prior to each measurement, excess NeutrAvidin was removed by flushing each chamber with ~1 mL buffer followed by incubation with lipid nanodisc for at least 5 min to ensure immobilisation. Excess nanodiscs were removed by flushing with buffer.

Acquisition of smFRET data

All smFRET experiments were performed on a total internal reflection fluorescence (TIRF) microscope (IX83, Olympus) equipped with two EMCCD cameras (imagEM X2, Hamamatsu) and an oil immersion 100x objective (UAPON 100XOTIRF, Olympus). Cy3 and Cy5 fluorophores were excited using 532 nm and 640 nm solid state laser lines (Olympus), respectively. A quad band filter cube was used to block the lasers in the emission pathway, while a multichannel imaging system (DC2 two-channel system, Photometrics) was used to split the signal into two channels. 582/75 nm and 700/75 nm band pass ET filters were used to filter donor (Cy3) and acceptor (Cy5) emission, respectively. All experiments were performed using ALEX as previously described (*30*, *33*) with 200 ms temporal resolution. All experiments were done in imaging buffer (50 mM TRIS, 600 mM NaCl, pH 7.9) with triplet quenchers (2 mM trolox, 2 mM para-nitrobenzyl alcohol, 2 mM cyclooctatetraene) and an oxygen scavenging system (1 U/mL PCD, 2.5 mM PCA) to avoid blinking and improve photostability of the fluorescent dyes.

Quantitative analysis of smFRET data

Quantitative image analysis was done using homemade software written in Python as described previously (*30*). Time-dependent signal and background traces were extracted for each colocalized Cy3/Cy5 pair and evaluated based on criteria such as signal/background ratio, noisiness, photobleaching and stoichiometry. Extracted traces were sorted to ensure only traces corresponding to single donor and acceptor fluorophores with anti-correlated signals were used for further analysis.

To correct FRET values for spectral crosstalk, direct excitation and detection efficiency we followed the method precisely outlined by Hellenkamp et al. (*53*) based on established methodology as first described by Lee et al. (*33*). In brief, we obtained α and δ correction factors for spectral crosstalk and direct excitation by manual inspection of the traces. Afterwards, we obtained the β and γ factors from FRET versus stoichiometry 2D histograms. The corrected FRET efficiencies were converted to distances using a Cy3/Cy5 Förster radius of 56 Å.

The distributions of FRET values for each experimental condition were fitted with a mixture of gaussians using unbinned likelihood fitting. Although FRET values are not technically Gaussian distributed, it has in practice been shown to be a robust method with little discrepancy (*17*, *29*, *30*). The number of underlying states (i.e. gaussians) was determined based on Bayesian Information Criterion (BIC). Since BIC is highly sensitive to the number of data points and each experimental condition had a different number of accepted FRET traces, the number of underlying states was determined from the largest data set as well as all data combined both resulting in five gaussians as the best model (Supplementary Fig 8). Hidden Markov modeling (HMM) was used to identify dynamic transitions as previously described.

#### Simulation of FRET traces and cross-correlation analysis

Traces were generated based on a two-state hidden markov model (HMM) with a transition probability of 0.95 between FRET states at 0.3 and 0.7, respectively. 8% gaussian noise was added to donor and acceptor intensities to simulate experimental uncertainty. To mimic various temporal resolutions, traces were binned at 1, 5, 10, 20, 50 and 100 frames per bin effectively causing between 0.95 and 95 transitions on average per bin at the various conditions. After binning, all traces were cut to a length of 200 frames with no bleaching to ensure equal amounts of data at different temporal resolutions. The Pearson cross-correlation was calculated for donor and acceptor intensities of each trace subsequent to bining.

#### Statistical analysis of *in vitro* activity data

The average and standard error of the mean (SEM) of at least three independent replicates were calculated followed by normalization to the average and SEM of at least three independent control measurements. All calculations were done in python with error propagation. P-values were calculated using Welch's t-test (unequal variances). Level of significance is marked by asterisk symbols (* $p<0.05$; ** $p<0.01$; *** $p<0.005$). All reported uncertainties are SEM unless otherwise stated.

#### Fitting of dose-response (IC50) curves

Dose-response curves from POR activity assays were fitted using the Hill equation in order to determine the IC50 value:

$$f(x) = A + \frac{B - A}{1 + \left(\frac{IC50}{x}\right)^{n}}$$

Here, A represents the starting point (no ligand), B represents the end point (maximum response) and n represents the Hill coefficient. The Hill coefficient was bounded to the interval [0,5] in the presence of rifampicin and cyclophosphamide. Dhurrin IC50 values were extracted from lower concentrations only (solid lines in Fig 3B) as the effect is inverted at concentrations above 100-500 μM dependent on the assay. This is not due to photophysics (Supplementary Fig 4). The activity loss at high dhurrin concentrations may indicate the presence of a feedback loop where high dhurrin concentrations inhibit POR electron transferring activity. This is especially important as POR is crucial for dhurrin formation and indicates a self-regulatory cue inhibiting POR function when high concentrations of dhurrin are produced. While the mechanism for this is unclear, it could originate from an alternative docking site with lower affinity. Deciphering this, however, extends beyond the scope of this work and is worth further investigation in future studies.

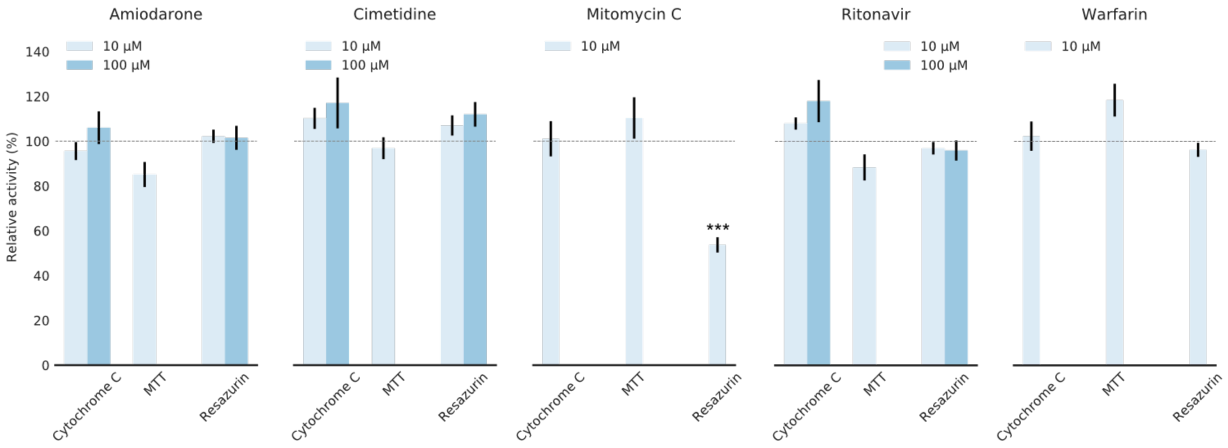


**Supplementary Fig. 1.** *In vitro* assays of small-molecule ligands showing weak or no effects on human POR proteoliposome activity. Barcharts display activity normalized to DMSO controls in the Cyt*c*, MTT and RS assay, respectively. Mitomycin C acts as a specific inhibitor/antagonist towards RS reduction (54±3 % of control). The remaining compounds have no statistically significant effect on POR function at the tested concentrations. Error bars represent ±SEM of at least three replicates with error propagation (see SI methods for details). The level of significance is marked by asterisk symbols (* $p<0.05$; ** $p<0.01$; *** $p<0.005$).

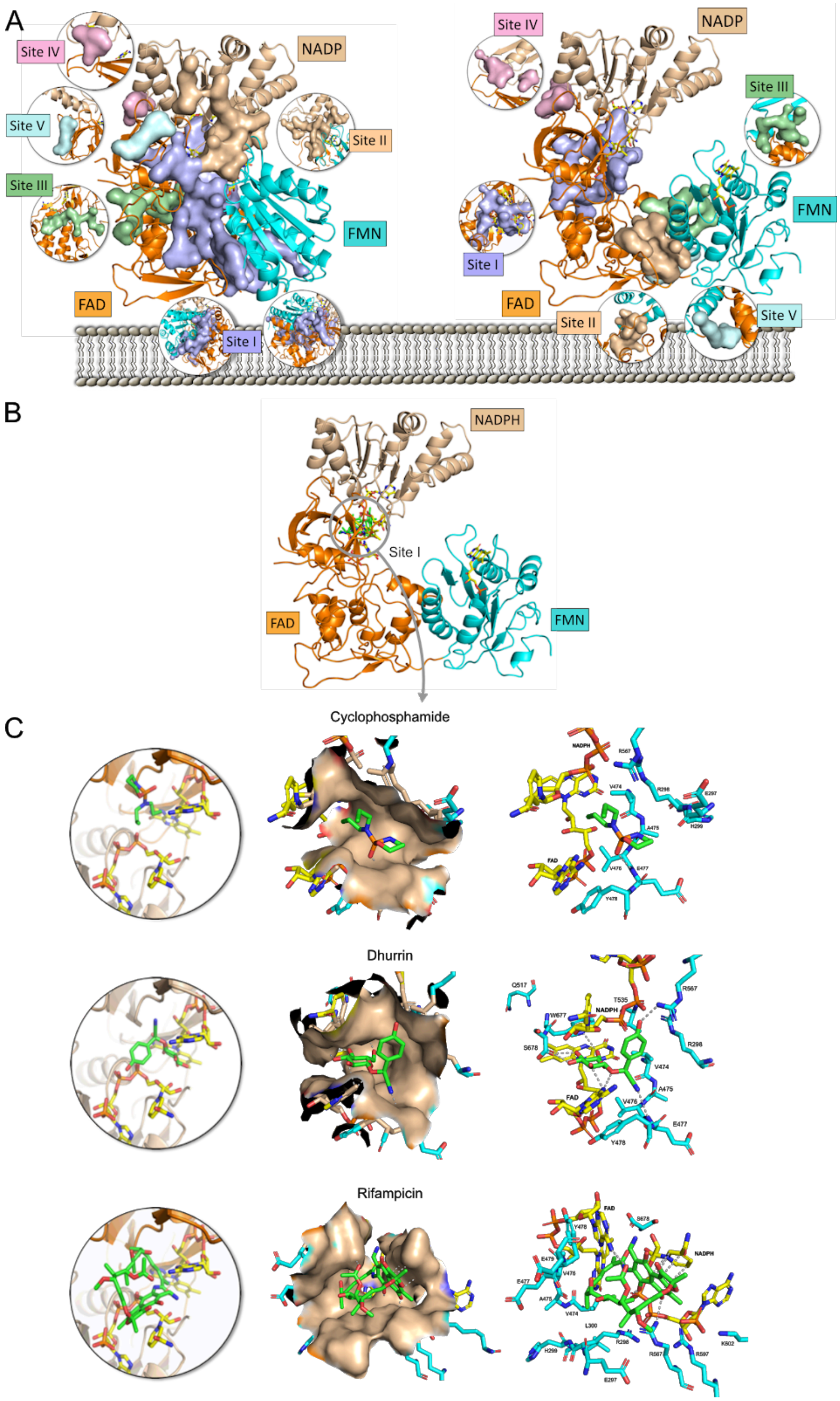


**Supplementary Fig. 2.** Potential ligand binding sites and docking of small-molecule ligands on rat POR. A) SiteMap analysis on human POR in a compact conformation (PDB 3QE2; left) and rat POR in an extended conformation (PDB 3ES; right) identifying five possible ligand binding sites (Sites I-V) on both isoforms. Sites I-III display SiteScore and Dscore values indicating that ligands may bind to these sites with sub-micromolar affinity (see Supplementary Table 1). Note, Sites I-V on human POR do not completely align with Sites I-V on rat POR. B) Docking of small-molecule ligands on rat POR in an extended conformation (PDB 3ES9). Ligands are displayed in green, while cofactors are displayed in yellow. C) Predicted docking of cyclophosphamide, dhurrin and rifampicin docking in Site I. All amino acid residues (blue) and cofactors (yellow) within 5 Å from the respective ligands are displayed. Cyclophosphamide is predicted to form H-bonds to A475 (2.8 Å) and E477 (3.1 Å). Dhurrin is predicted to form H-bonds to E477 (3.5 Å), R567 (2.8 Å), W677 (3.3 Å), S678 (2.7 Å and 2.8 Å), NADPH (2.6 Å) and FAD (2.9 Å, 3.0 Å, 3.1 Å). Rifampicin is predicted to form H-bonds to R567 (3.0 Å), NADPH (2.9 Å and 3.0 Å) and FAD (3.3 Å), and pi-pi interactions with NADPH (3.7-4.3 Å). See Supplementary Table 1 for predicted binding energies.

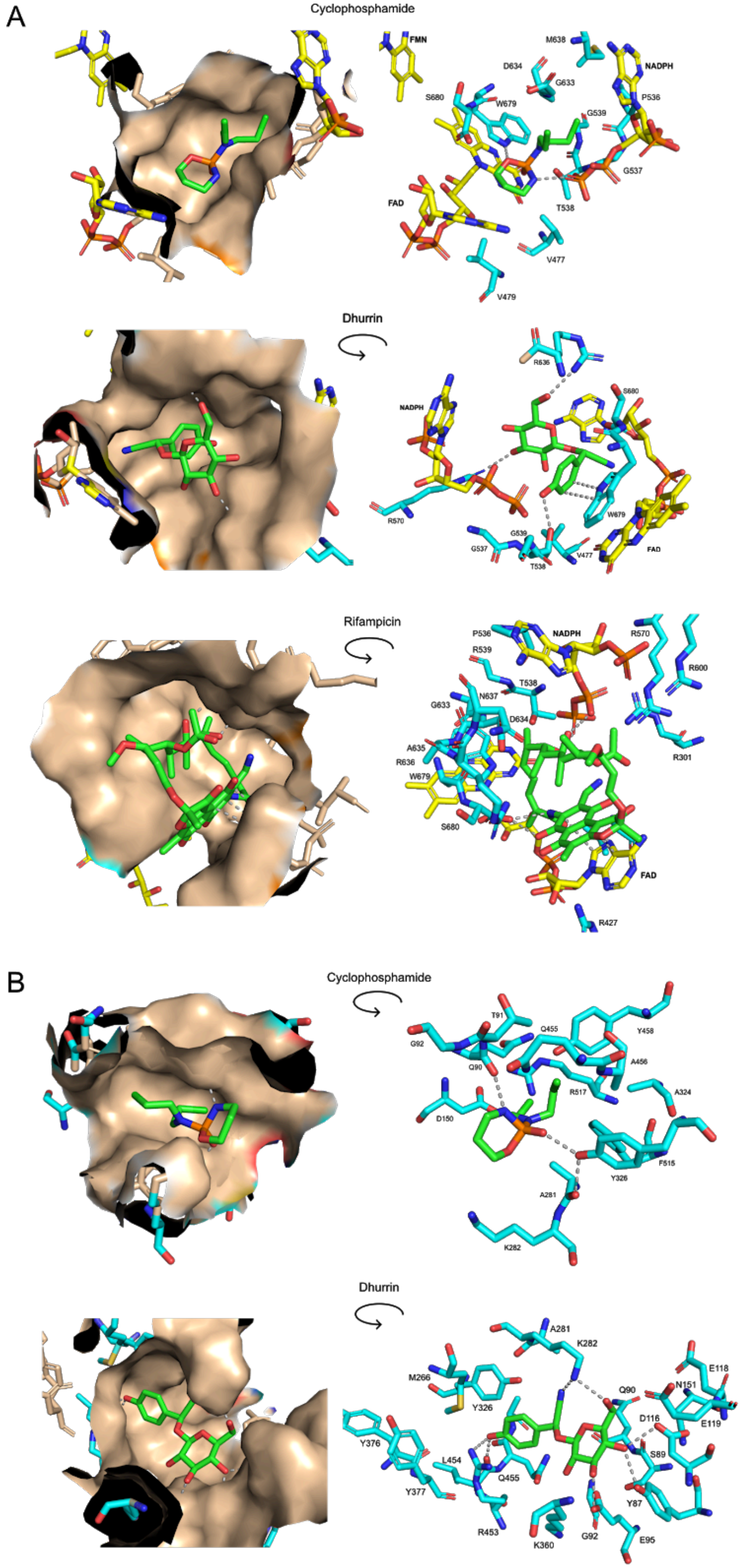


**Supplementary Fig. 3.** Predicted docking conformations of small-molecule ligands on human POR (PDB 3QE2) in Site Ia (A) and Site Ib (B). All amino acid residues (blue) and cofactors (yellow) within 5 Å from the respective ligands (green) are displayed. In Site Ia, Cyclophosphamide appears to dock in a cavity with the chlorines tightly fitting into a groove partly formed by NADPH. It is predicted to form H-bonds to W679 (3.3 Å) and NADPH (2.6 Å). Dhurrin also docks in a cavity between NADPH and FAD with predicted H-bonds to T538 (2.9 Å), R570 (3.2 Å), R636 (2.9 Å) and pi-pi interactions with W679 (3.6-4.2 Å). Rifampicin is predicted to form pi-pi interactions with FAD (3.6-4.3 Å), a H-bond to S680 (2.8 Å) and two H-bonds to NADPH (2.4 Å and 3.2 Å). Notably, amino acid residues G539 and R600 which are within 5 Å from the three ligands in Site Ia are both associated with POR deficiency. Pathogenic mutations G539R and R600W both cause disorder of sexual development due to low production of sex steroids (*5*, *14*). In Site Ib, cyclophosphamide is predicted to form H-bonds to A475 (2.8 Å) and E477 (3.1 Å). Dhurrin is predicted to form H-bonds to E477 (3.5 Å), R567 (2.8 Å), W677 (3.3 Å), S678 (2.7 Å and 2.8 Å), NADPH (2.6 Å) and FAD (2.9 Å, 3.0 Å, 3.1 Å), and rifampicin is predicted to form H-bonds to R567 (3.0 Å), NADPH (2.9 Å and 3.0 Å) and FAD (3.3 Å), and pi-pi interactions with NADPH (3.7-4.3 Å).

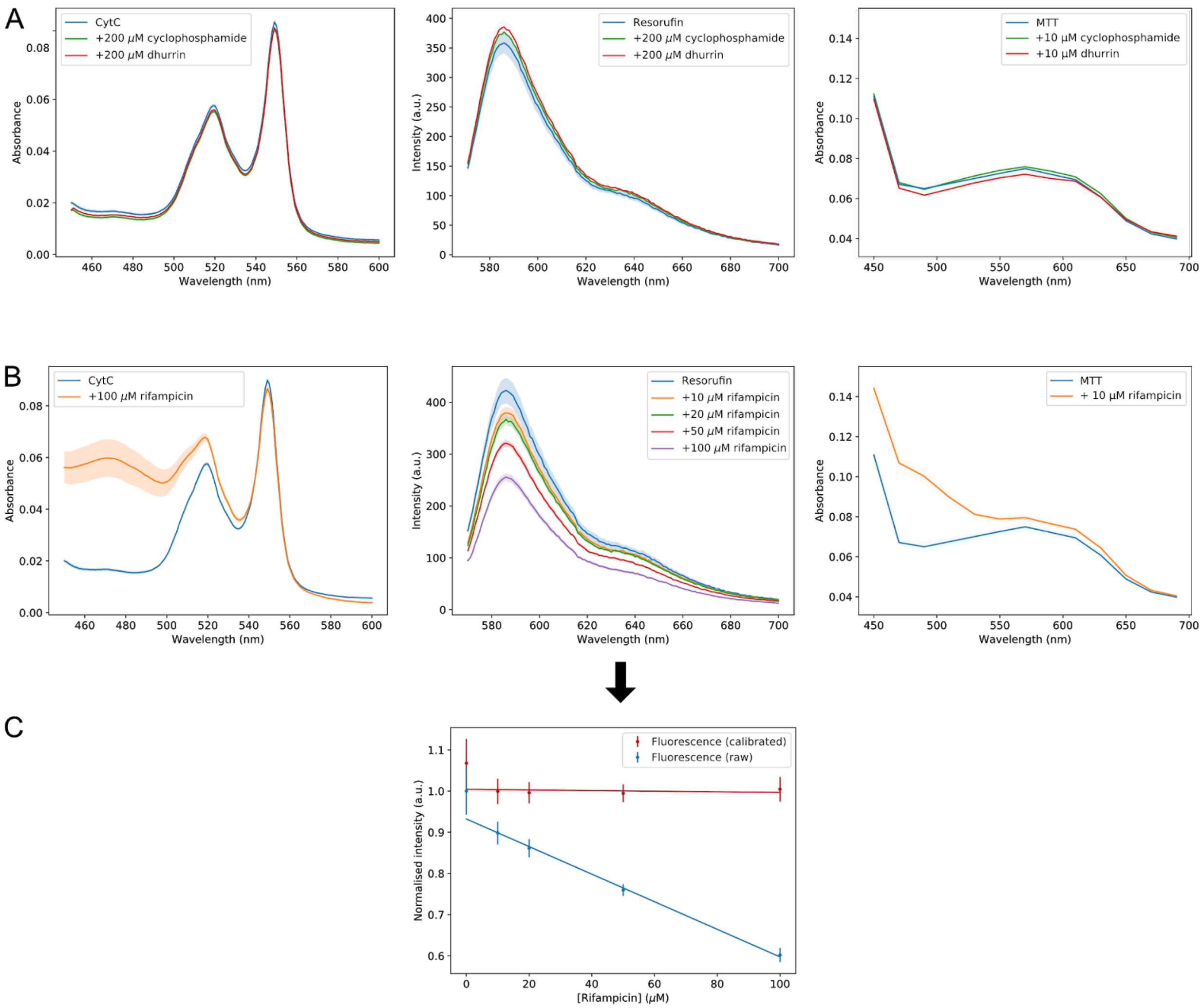


**Supplementary Fig. 4.** Control spectra and calibration of small-molecule ligands on Cytc, resorufin and MTT spectral properties. A) Control spectra showing that neither cyclophosphamide nor dhurrin induce any photophysical effects on Cyt*c* absorbance, resorufin emission nor MTT absorbance. B) Control spectra of rifampicin on Cyt*c* absorbance, resorufin emission (540 nm excitation) and MTT absorbance. Rifampicin does not significantly affect the absorbance readout of Cyt*c* nor MTT at the relevant wavelengths (550 nm and 610 nm, respectively), however resorufin emission is quenched by rifampicin in a concentration dependent manner (10-100 μM rifampicin). C) Calibration curve showing a linear correlation between rifampicin concentration and fluorescence intensity of resorufin extracted at the 585 nm emission peak (blue curve). To account for fluorescence quenching in the presence of rifampicin, a calibration was performed (red curve). Error bars represent ±SEM of triplicate measurements. All rifampicin measurements presented in this study have been calibrated accordingly (RS assay only). A-B) Each spectrum represents the mean ± SEM of triplicate measurements (shaded area).

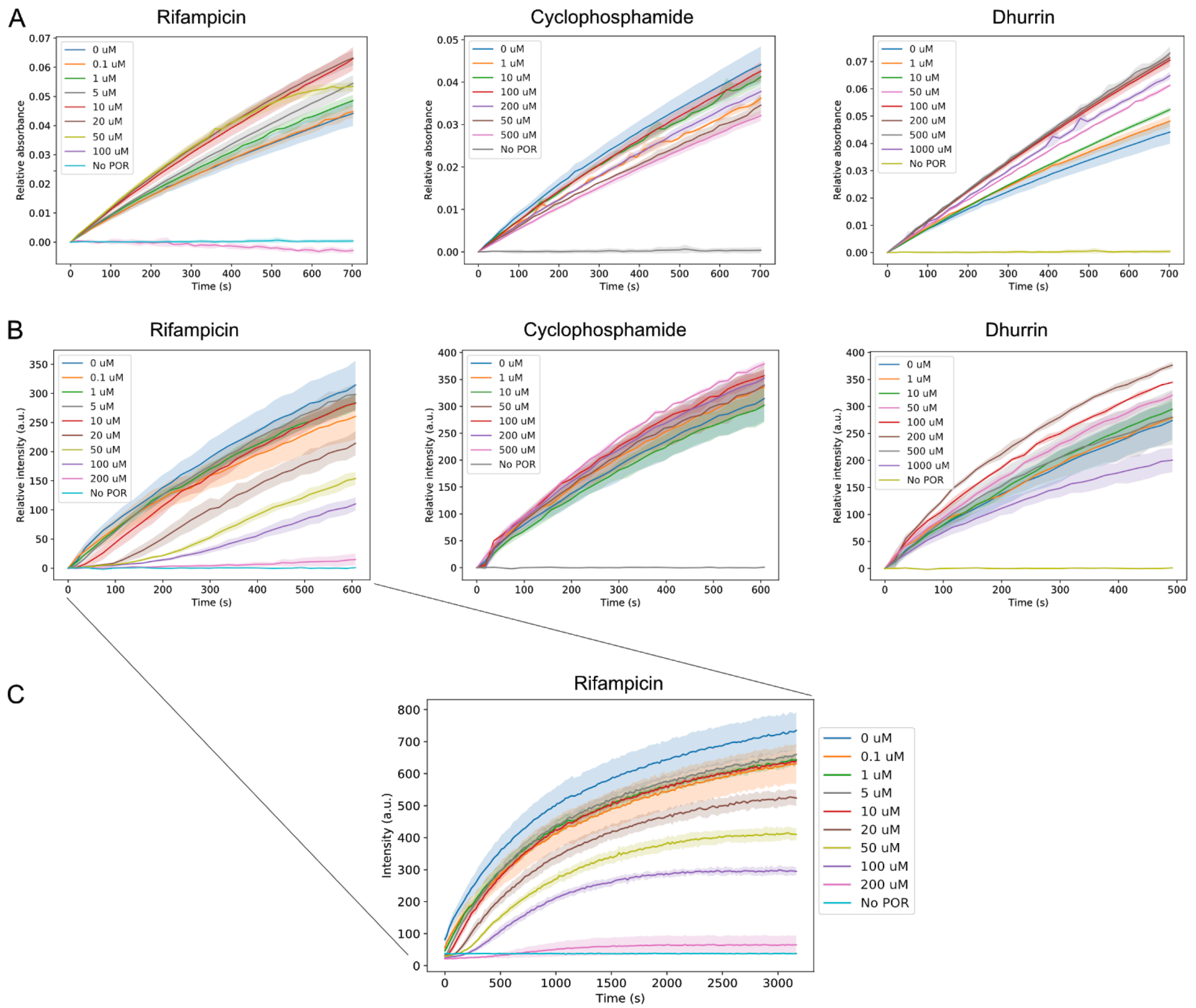


**Supplementary Fig. 5.** Raw activity traces of an experiment on *Sb*POR2b proteoliposomes in A) Cyt*c* assay and B) RS assay. The linear region of each trace was used to quantify activity. Rifampicin traces are calibrated according to Supplementary Fig 4 and the activity is extracted after the lag phase (discussed in panel C). Each trace represents the mean ± SD of at least triplicate measurements (shaded area). C) Rifampicin induced a lag phase in *Sb*POR2b proteoliposome activity towards RS. Time trajectories of emission intensity are shown in the presence of varying rifampicin concentrations (0-200 µM). Reaction rates were extracted after the lag phase for all screening and dose-response experiments. Each trace represents the mean ± SD of triplicate measurements (shaded area). We note that the lag phase is only observed for the *Sb*POR2b isoform towards RS. It is not apparent using human POR in neither detergent micelles nor proteoliposomes. Deciphering the mechanism underlying the lag phase extends beyond the scope of this work, however, is worth investigation in future studies.

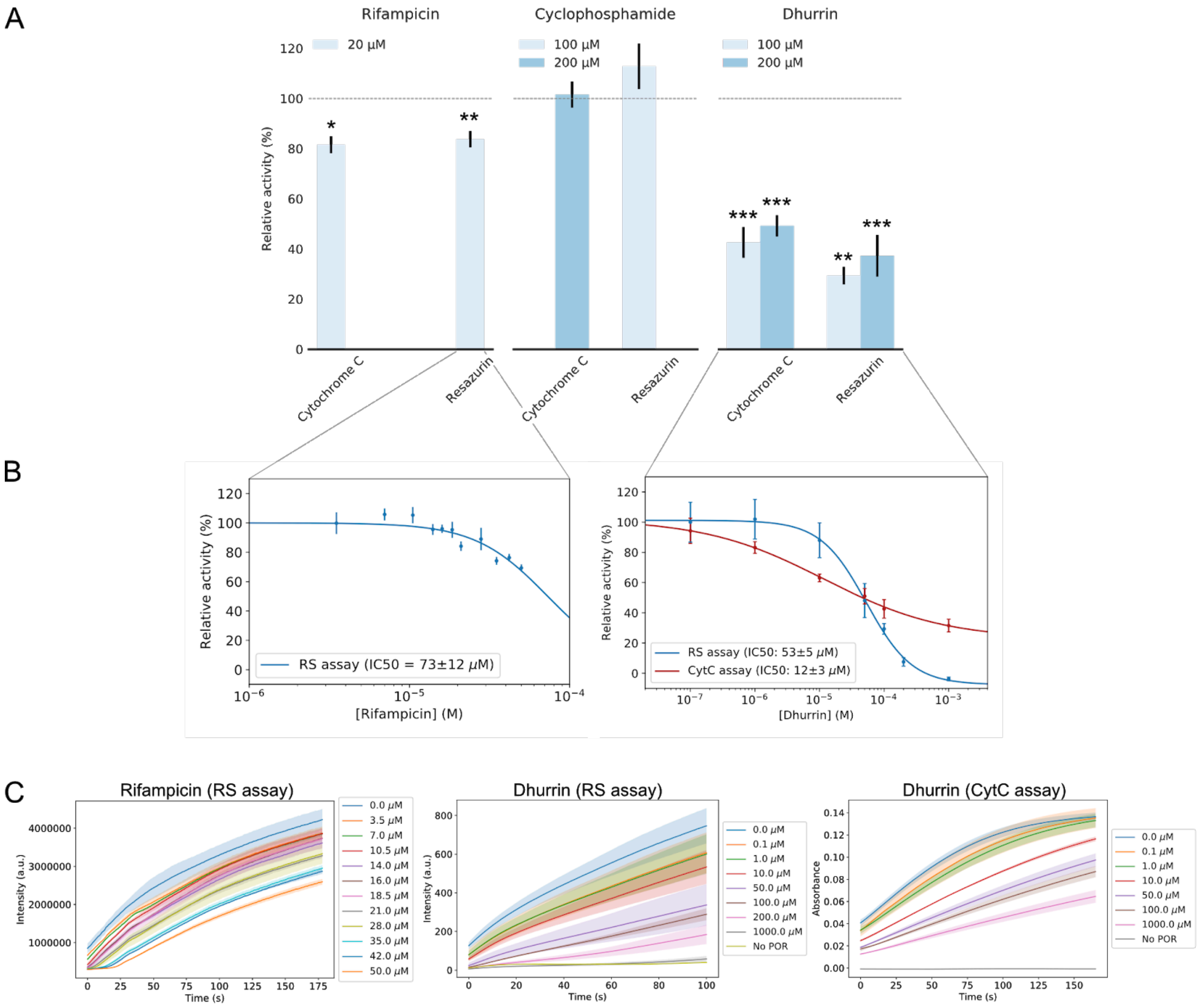


**Supplementary Fig. 6.** *In vitro* assays of small-molecule ligands on *Sb*POR2b in detergent micelles. A) Bar chart displaying POR activity normalized to controls in the Cyt*c* and RS assay, respectively. Level of significance is marked by asterisk symbols (* p<0.05; ** p<0.01; *** p<0.005). B) Dose-response curves of rifampicin (left) and dhurrin (right). IC50 values are depicted in the figure legend (low micromolar range). C) Raw activity traces of *Sb*POR2b underlying the dose-response curves. The linear region of each trace was used to quantify activity. For rifampicin traces, the activity was extracted after the lag phase according to Supplementary Fig 5. Each trace represents the mean ± SD of at least triplicate measurements (shaded area). A-B) Error bars represent ±SEM from at least three replicates with error propagation.

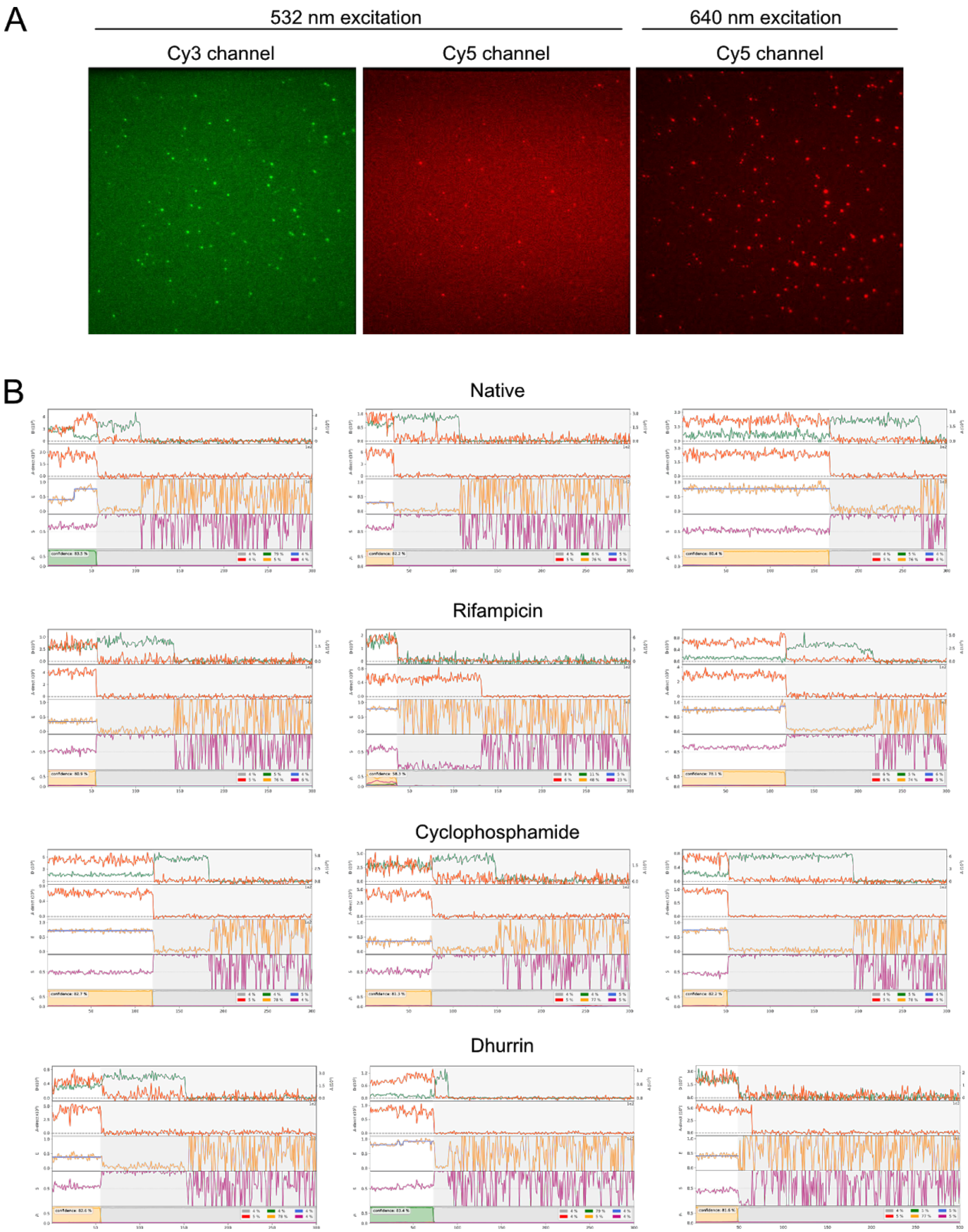


**Supplementary Fig. 7.** Raw TIRF microscope images and representative smFRET traces. A) TIRF microscope images displaying one field-of-view (82x82 µm) of a representative surface with diffraction-limited, immobilized, dual-labelled *Sb*POR2b in nanodiscs excited at 532 nm and 640 nm, respectively. The images are acquired using emission filters optimized for Cy3 and Cy5 fluorophores (see SI methods) and show one frame from movies recorded at 200 ms temporal resolution. B) Representative smFRET traces at each experimental condition (A-D). Every trace has four panels: The top panel displays donor (green) and acceptor (red) intensities at 532 nm excitation. The second panel displays acceptor intensity (red) at 640 nm excitation using ALEX.

The third panel displays the FRET value (orange) calculated with calibration factors, and idealized FRET value determined from HMM fitting (blue). The fourth panel displays the donor-acceptor stoichiometry, which should ideally be 0.5 at a 1:1 donor-acceptor ratio. Some of the depicted traces display a static FRET value while others display transitions between long-lived equilibrium states. The x-axis represents frames (frame rate: 5 $s^{-1}$).

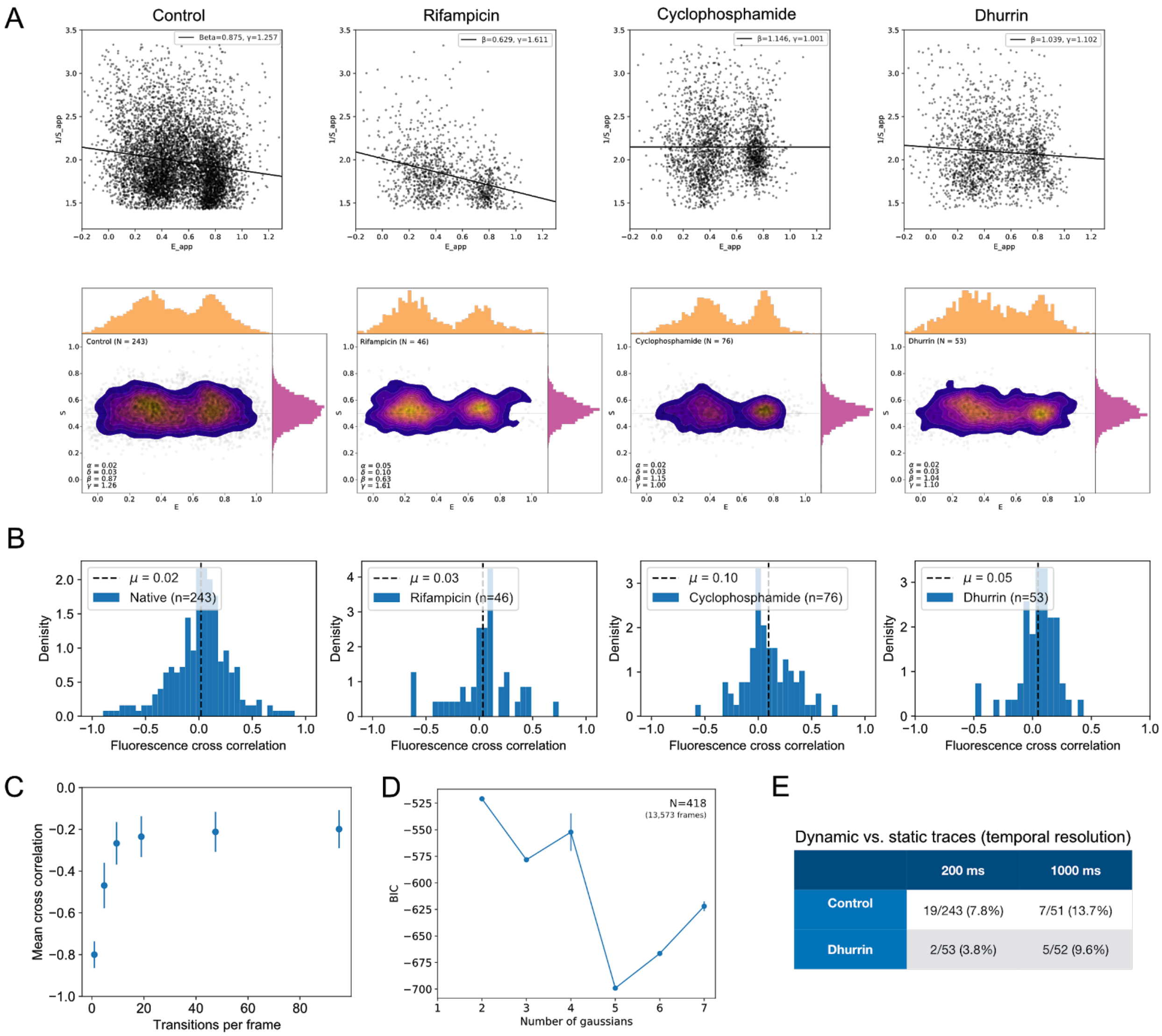


**Supplementary Fig. 8.** Correction factors, fluorescence cross correlation and BIC analysis of smFRET data. A) Determination of β and γ correction factors from global fitting of ALEX smFRET data following published methodology (*53*) (top), and 2D histograms displaying smFRET data after applying correction factors (bottom). B) Pearson correlation histograms of *Sb*POR2b smFRET data in the native form and in the presence of rifampicin, cyclophosphamide and dhurrin, respectively, displaying no autocorrelation in agreement with recent published data (*7*, *26*). C) Simulation of smFRET traces displaying the mean cross correlation of simulated donor and acceptor intensities for varying temporal resolution in agreement with earlier studies of fast conformational transitions on GPCRs (*17*). A two-state kinetic model with a 95% transition probability between each conformational state was used to simulate donor and acceptor trajectories without bleaching. 8% gaussian noise was added to each trace to mimic experimental uncertainty. The traces were subsequently binned using bin sizes varying from 1 to 100 to mimic various temporal resolutions. Increasing the bin size (i.e. average number of transitions per frame) results in loss of anticorrelation. Data display mean ± SD of 1024 simulated traces. D) Bayesian Information Criterion (BIC) scores of pooled *Sb*POR2b smFRET data (418 particles) from fitting gaussian mixture models ranging from 2 to 7 gaussians. The lowest BIC score is obtained with a 5-state gaussian mixture model. Error bars represent ±SEM from bootstrapping (n=20). E) Fraction of smFRET traces showing dynamic transitions as a function of temporal resolution. The fraction of traces showing transitions increases slightly when decreasing temporal resolution from 200 ms to 1 s, indicating that traces do not display an ensemble average but rather transitions between long-lived equilibrium states. The increase in observed transitions at 1 s as compared to 200 ms temporal resolution is caused by longer observation times allowing for more transitions to occur before photobleaching.

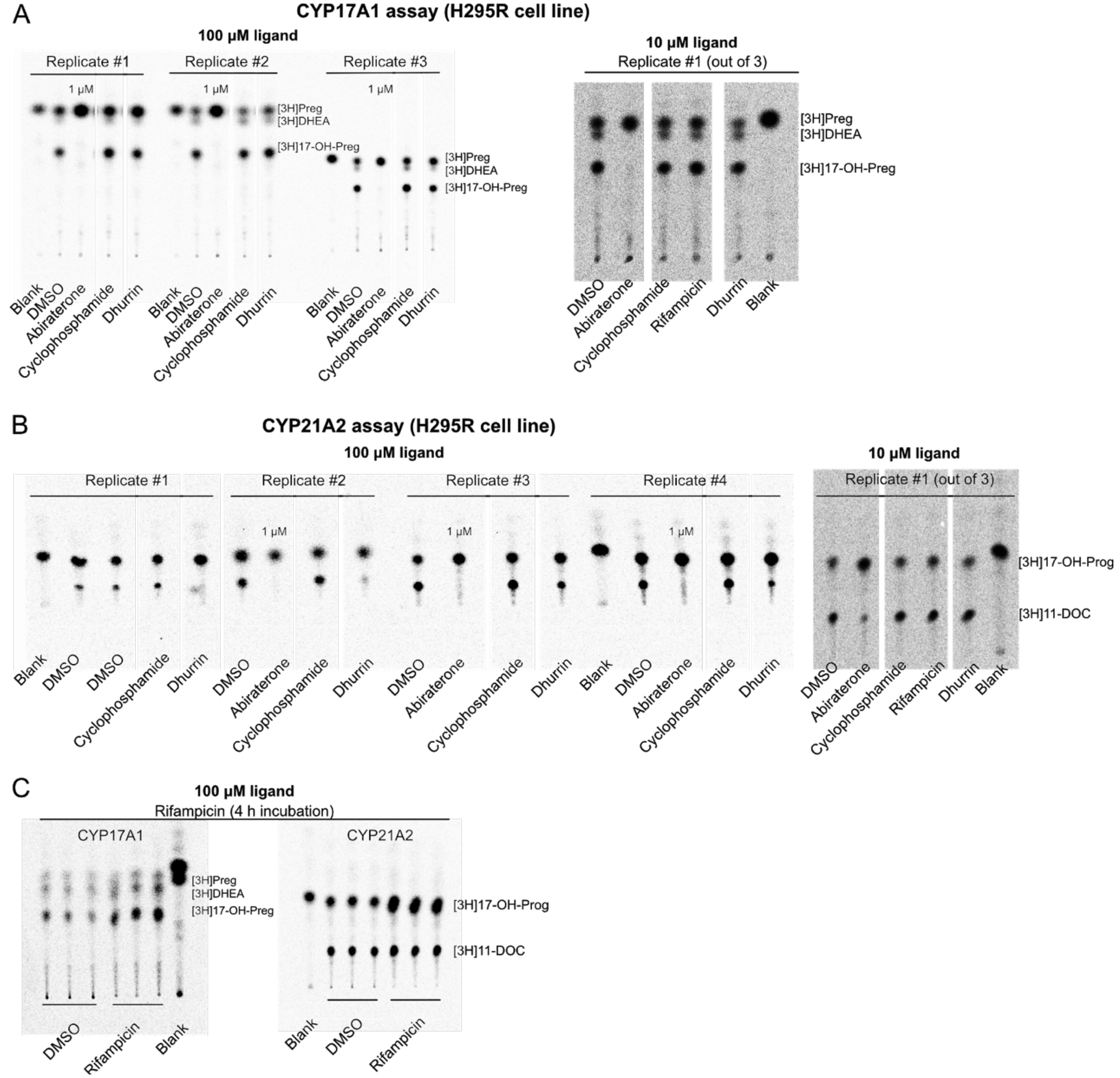


**Supplementary Fig. 9.** Raw TLC images of CYP17A1 activity assay (A) and CYP21A2 activity assay (B) in H295R cells at 100 µM and 10 µM ligand concentrations. Cells were incubated with ligand for 24 h before adding radiolabeled substrate and probing activity. Radiolabeled substrates and products were visualized on a PhosphorImager and quantified using Multi Gauge software. C) CYP17A1 and CYP21A2 assay at 100 µM rifampicin upon 4 h of incubation.

| Protein | Site | SiteScore | Dscore | Volume (Å3) | Residues defining the site |
|---|---|---|---|---|---|
| 3ES9 | Site II | 1.06 | 0.98 | 584 | 97,101,104,105,214,217,218,220,221,224,225,228,235,240,241,242,243,244,245,357,360,361,375,381,382,384,385,387,388,446,447,448,449,450 |
| | Site I | 1.00 | 0.96 | 751 | 291,297,298,299,325,326,327,328,330,333,378,379,424,426,428,429,432,452,453,454,455,474,475,476,477,478,484,485,486,487,488,489,492,493,495,496,533,534,535,536,567,631,632,634,636,677,678,752,753 |
| | Site III | 0.93 | 0.95 | 273 | 198,199,200,201,202,203,204,205,215,218,219,222,225,226,229,230,382,383,404,406,407,408,409,410,411,412,413,416,417 |
| | Site IV | 0.84 | 0.74 | 158 | 290,291,292,293,294,302,304,470,541,545,569,570,571,572,573,574,575,576,577,578 |
| | Site V | 0.71 | 0.65 | 128 | 228,229,230,232,233,234,235,240,241,387,388,390,391,403 |
| 3QE2 | Site I | 1.03 | 0.98 | 2988 | 69,75,78,79,81,87,89,90,91,92,93,95,96,97,99,100,102,103,104,105,106,107,108,111,112,113,114,115,116,118,119,120,143,149,150,151,178,181,211,212,213,214,215,216,220,223,224,227,228,231,237,238,243,244,245,246,247,248,262,266,267,278,279,280,281,282,294,299,300,301,302,313,314,315,316,317,318,326,353,354,355,357,358,359,360,361,362,363,364,365,366,378,379,380,381,382,383,384,385,386,387,388,390,391,409,410,411,412,419,424,427,430,448,449,450,451,452,453,454,455,456,457,458,477,478,479,480,481,494,498,515,516,517,518,536,537,538,539,570,633,634,636,638,679,680,751,752,753 |
| | Site II | 1.00 | 0.98 | 689 | 90,143,144,145,146,147,148,149,150,153,179,180,181,182,183,184,318,319,322,458,466,516,517,518,519,520,521,522,523,524,525,526,527,546,549,550,553,628,630,631,635,636,637,639,640,642,643,647,666,669,670,672,674,675,676,677,678,751,752 |
| | Site III | 1.00 | 1.02 | 456 | 263,264,265,270,272,273,286,328,329,330,331,332,333,334,336,371,375,376,380,381,382,383,427,428,429,431,432,435,457,479,481,483,487,488,489,490,491,492,495,496,499,500,509,510,511,513,752 |
| | Site V | 0.65 | 0.49 | 74 | 310,311,312,313,314,315,316,317,318,465,469,470 |
| | Site IV | 0.60 | 0.53 | 78 | 293,294,295,296,297,572,574,575,576,577,578,579,580,581 |

| Protein | Ligand | Site | Gscore | Emodel | Residues within 5 Å from the ligand |
|---|---|---|---|---|---|
| 3ES9 | Cyclophosphamide | Site I | -5.0 | -40.5 | E297, R298, H299, A475, V476, E477, Y478, R567, FAD, NADPH |
| | | Site II | -4.6 | -32.2 | - |
| | | Site V | -4.4 | -36.7 | - |
| | | Site IV | -4.2 | -30.6 | - |
| | | Site III | -4.1 | -32.3 | - |
| | Dhurrin | Site I | -7.0 | -67.4 | R298, V474, A475, V476, E477, Y478, T535, R567, W677, S678 |
| | | Site II | -5.7 | -56.8 | - |
| | | Site V | -5.2 | -54.3 | - |
| | | Site III | -5.1 | -47.8 | - |
| | | Site IV | -4.7 | -44.9 | - |
| | Rifampicin | Site I | -4.6 | -51.2 | E297-L300 (ERHL), V474-E479 (VAVEYE), R567, R597, K602, S678 |
| 3QE2 | Cyclophosphamide | Site Ia | -5.2 | -45.3 | V477, V479, P536-**G539*** (PGTG), G633, D634, M638, W679, S680, FAD, NADPH |
| | | Site Ib | -5.0 | -45.3 | Q90, T91, Q92, D150, A281, K282, A324, Y326, Q455, A456, Y458, F515, R517 |
| | | Site III | -4.5 | -35.7 | - |
| | | Site IV | -4.1 | -31.7 | - |
| | | Site II | -3.9 | -34.2 | - |
| | | Site V | -3.0 | -22.4 | - |
| | Dhurrin | Site Ib | -6.7 | -68.3 | Y87, S89, Q90, G92, E95, E118, E119, N151, M266, A281, K282, Y326, K360, Y376, Y377, R453, L454, Q455 |
| | | Site Ia | -6.3 | -69.0 | V477, G537, T538, **G539***, R570, G633, D634, R636, W679, S680, FAD, NADPH |
| | | Site III | -6.2 | -67.1 | - |
| | | Site II | -5.7 | -60.5 | - |
| | | Site V | -5.6 | -47.3 | - |
| | | Site IV | -5.3 | -46.6 | - |
| | Rifampicin | Site Ia | -6.1 | -83.6 | R301, R427, V479, P536, T538, **G539***, R570, **R600***, G633-N637 (GDARN), W679, S680, FAD, NADPH |

* Residues associated with POR deficiency

**Supplementary Table 1.** Top) Potential ligand binding sites on human POR in a compact conformation (PDB 3QE2) and rat POR in an extended conformation (3ES9) identified from SiteMap analysis. Sites I-III display SiteScore and Dscore values indicating that ligands may bind to these sites with submicromolar affinity. Note, Sites I-V on human POR do not completely align with Sites I-V on rat POR. Bottom) Binding energies (Gscore; kcal/mol) and Emodel scores of small-molecule ligands on human POR in a compact conformation (PDB 3QE2) and rat POR in an extended conformation (PDB 3ES9) predicted from computational docking simulations. Only the lowest energy score of each ligand in each site is considered. Amino acid residues within 5 Å from the docked ligands in Sites Ia and Ib of human POR and Site I of rat POR are shown. Human POR residues associated with POR deficiency are marked as bold. G539R and R600W are both found in patients with POR deficiency and cause disorder of sexual development due to low production of sex steroids (*5*, *14*).
